# Isoform-specific characterization implicates alternative splicing in *APOBEC3B* as a mechanism restricting APOBEC-mediated mutagenesis

**DOI:** 10.1101/2020.09.27.315689

**Authors:** A. Rouf Banday, Olusegun O. Onabajo, Seraph Han-Yin Lin, Adeola Obajemu, Joselin M. Vargas, Krista A. Delviks-Frankenberry, Philippe Lamy, Ariunaa Bayanjargal, Clara Zettelmeyer, Oscar Florez-Vargas, Vinay K. Pathak, Lars Dyrskjøt, Ludmila Prokunina-Olsson

## Abstract

APOBEC3A (A3A) and APOBEC3B (A3B) enzymes drive APOBEC-mediated mutagenesis, but the understanding of the regulation of their mutagenic activity remains limited. Here, we showed that mutagenic and non-mutagenic A3A and A3B enzymes are produced by canonical and alternatively spliced *A3A* and *A3B* isoforms, respectively. Notably, increased expression of the canonical *A3B* isoform, which encodes the mutagenic A3B enzyme, predicted shorter progression-free survival of bladder cancer patients. Expression of the mutagenic *A3B* isoform was reduced by exon 5 skipping, generating a non-mutagenic *A3B* isoform. The exon 5 skipping, which was dependent on the interaction between SF3B1 splicing factor and weak branch point sites in intron 4, could be enhanced by an SF3B1 inhibitor, decreasing the production of the mutagenic A3B enzyme. Thus, our results underscore the role of A3B, especially in bladder cancer, and implicate alternative splicing of *A3B* as a mechanism and therapeutic target to restrict APOBEC-mediated mutagenesis.

## INTRODUCTION

Enrichment of C-to-T or C-to-G mutations within the T<u>C</u>A and T<u>C</u>T motifs, attributed to APOBEC-mediated mutagenesis, has been implicated in cancer susceptibility^1^, tumor evolution^2,3^, metastatic progression^4,5^, treatment response^3,6^, and survival^1^. Thus, understanding and harnessing the mechanisms regulating this mutational process is of clinical importance. Among the seven members of the APOBEC3 enzyme family, A3A^7,8^, A3B^9^, and an allelic variant of APOBEC3H (A3H)^10^ have been linked with APOBEC mutagenesis in tumors.

The intrinsic and extrinsic factors that regulate the expression levels of *APOBEC3s* might explain some of the differences in the load of APOBEC-signature mutations within and between tumors of different types. These intrinsic factors include common germline variants - a single nucleotide polymorphism (SNP) rs1014971 and the *A3AB* deletion^1,11^, or their correlated proxies, SNPs rs17000526 and rs12628403^1^. The same genetic variants have also been associated with *A3B* expression, such as rs1014971 regulating *A3B* expression through an allele-specific effect on an enhancer upstream of the *APOBEC3* gene cluster^1^, and the *A3AB* deletion – through the elimination of one or both copies of the *A3B* gene^11,12^. A haplotype represented by a missense SNP rs139297 (Gly105Arg) that creates an A3H protein isoform with nuclear localization (A3H-I), has been associated with APOBEC-signature mutations in *A3A* and *A3B*-null breast and lung tumors^10^. Extrinsic factors that induce expression of specific *APOBEC3s* include viral infections^1,13^ and exposure to environmental or chemotherapeutic DNA-damaging agents^1,2,14^. Most of the conclusions about the role of A3A and A3B in mutagenesis were drawn from studies based onRNA-seq, despite poor ability to resolve and confidently quantify mRNA expression of these highly homologous APOBECs by short sequencing reads^10^.

Alternative splicing (AS) of pre-mRNA can produce functionally distinct isoforms or regulate gene expression through downstream mechanisms, such as nonsense-mediated decay (NMD)^15^. Alternatively spliced isoforms of other *APOBEC3* genes (*APOBEC3H* and *APOBEC3F*) have been reported to generate enzymes with variable activity^16-18^, but the effects of AS in *A3A* and *A3B* on functional activities of these enzymes and relevant clinical outcomes have not been explored. Here, we characterized AS in *A3A* and *A3B* in relation to APOBEC-mediated mutagenesis and explored the mechanisms of its regulation and possible therapeutic modulation.

## RESULTS

### Expression profiling based on specific exon-exon junctions provides more accurate quantification of *A3A* and *A3B* transcripts compared to total gene expression analysis by RNA-seq

According to the human reference genome annotation (hg19, UCSC), *A3A* and *A3B* genes have 2 and 3 alternative isoforms, respectively (**Table S1, Figure S1A**). Among these, only canonical isoforms, which we designated as *A3A1* and *A3B1*, but not alternatively spliced isoforms (*A3A2, A3B2*, and *A3B3)*, have been studied. First, we confirmed the existence of all these isoforms by analyzing RNA-seq reads in TCGA dataset and identifying exon-exon junction reads specific to each of these isoforms (**Figure 1A, Figure S1B**). The alternative *A3A* and *A3B* isoforms were expressed in 3.49% and >50% samples, respectively (**Figure 1B, C)**. Notably, we found quantification based on specific exon-exon junctions (by RNA-seq or qRT-PCR) more reliable than based on total gene expression by RNA-seq (**Note S1**).

**Figure 1.**
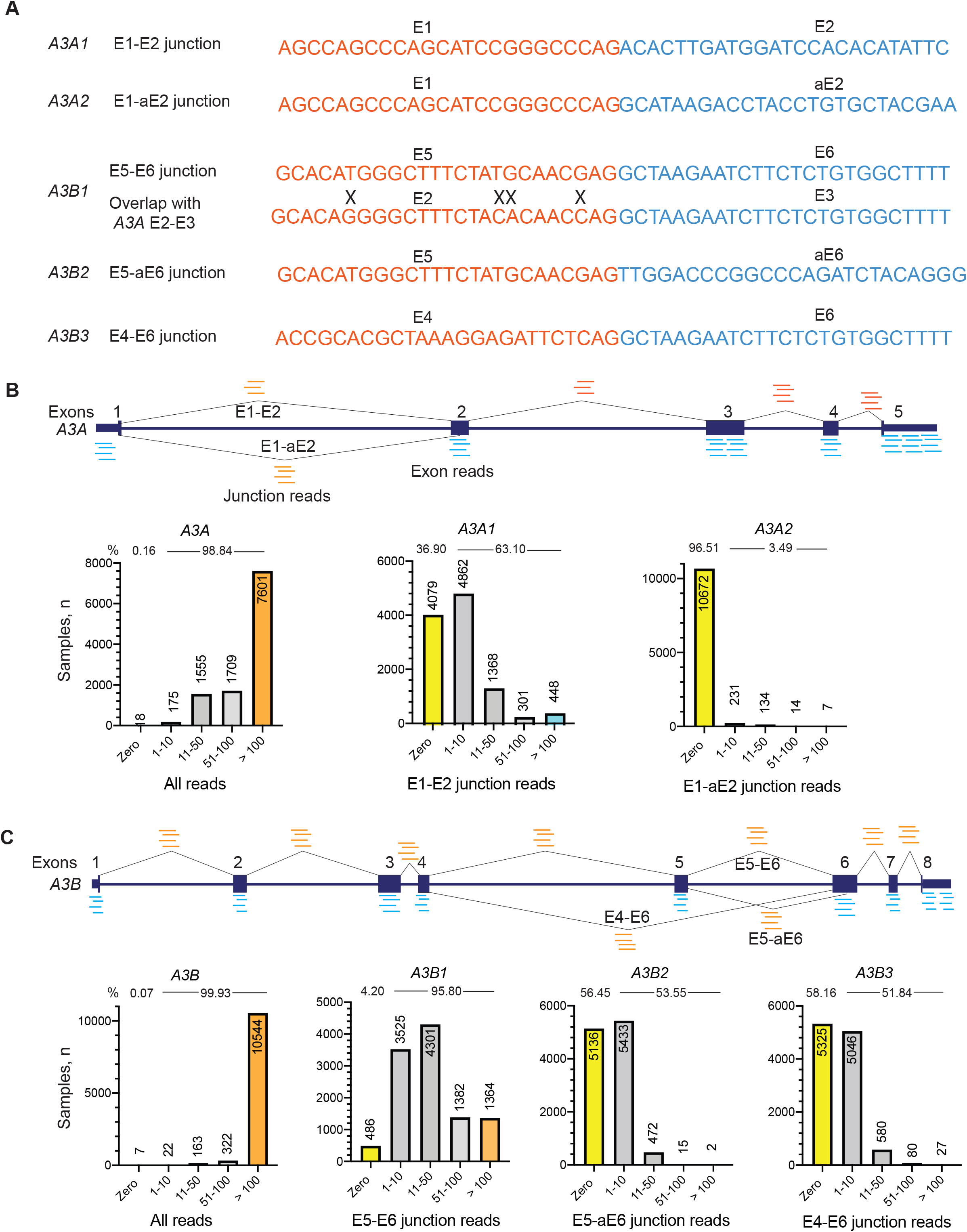
Quantification of *A3A* and *A3B* expression based on the total and exon-exon junction RNA-seq reads in 11,058 TCGA samples. **A)** Nucleotide sequences of exon-exon junctions specific to *A3A* and *A3B* isoforms. “X” – mismatch between *A3A* and *A3B* sequences. **B)** Schematics of *A3A* exons and splicing junctions. The bar graphs show the numbers of samples (y-axis) in relation to RNA-seq read counts grouped in 5 sub-categories (x-axis) for the total *A3A* expression and exon-exon junction-based expression of *A3A1* and *A3A2* isoforms. Based on exon junction reads (≥1 read/sample), *A3A1* (E1-E2 junction) is not expressed in most TCGA samples and *A3A2* is expressed only in 386 samples (3.49%). **C**) Schematics of *A3B* exons and splicing junctions and comparison of gene expression based on the total and exon-exon junction reads corresponding to *A3B* isoforms. *A3B2* and *A3B3* are expressed in 53.55% (5,922 of 11,058) and 51.84% (5,733 of 11,058) of TCGA samples, respectively. The TCGA RNA-seq set includes 10,328 tumors and 730 adjacent normal tissue samples.

This was also reflected by the analysis of *A3A* and *A3B* expression in TCGA samples comparing detection based on all gene-specific reads vs. exon-exon junction reads for specific isoforms. Based on total RNA-seq reads, *A3A* was undetectable only in 0.16% (18 of 11,058) of TCGA samples (**Figure 1B**). However, based on junction reads for the canonical isoform *A3A1 (*E1-E2 junction), expression was undetectable in 36.9% (4079 of 11,058) of TCGA samples (**Figure 1B**). Similarly, *A3B* expression was undetectable by total RNA-seq reads only in 0.07% (8 of 11,058) of TCGA samples, but this number was 4.2% (468 of 11,058) based on junction reads for the canonical *A3B1* isoform (E5-E6 junction reads, **Figure 1C**). Thus, due to high homology between *A3A* and *A3B* (**Note S1**), the misaligned RNA-seq reads would incorrectly represent the expression of both genes, affecting downstream analyses and biological interpretation.

### Alternative protein isoforms of A3A and A3B are non-mutagenic

To better understand the functional properties of these alternative isoforms, we performed computational analysis of their protein sequences. Compared to A3A1, A3A2 lacks 18 aa (residues 10-28) due to AS between exons 1 and 2, including residues His11 and His16, which are important for deamination activity^19^ (**Figure 2A**). A3B2 is produced due to AS in exon 6, resulting in the loss of 25 aa (residues 242-266, **Figure 2B**), including His253 that stabilizes the zinc cofactor, and Glu255 that directly participates in a nucleophilic attack on cytosine during deamination^20^.

**Figure 2.**
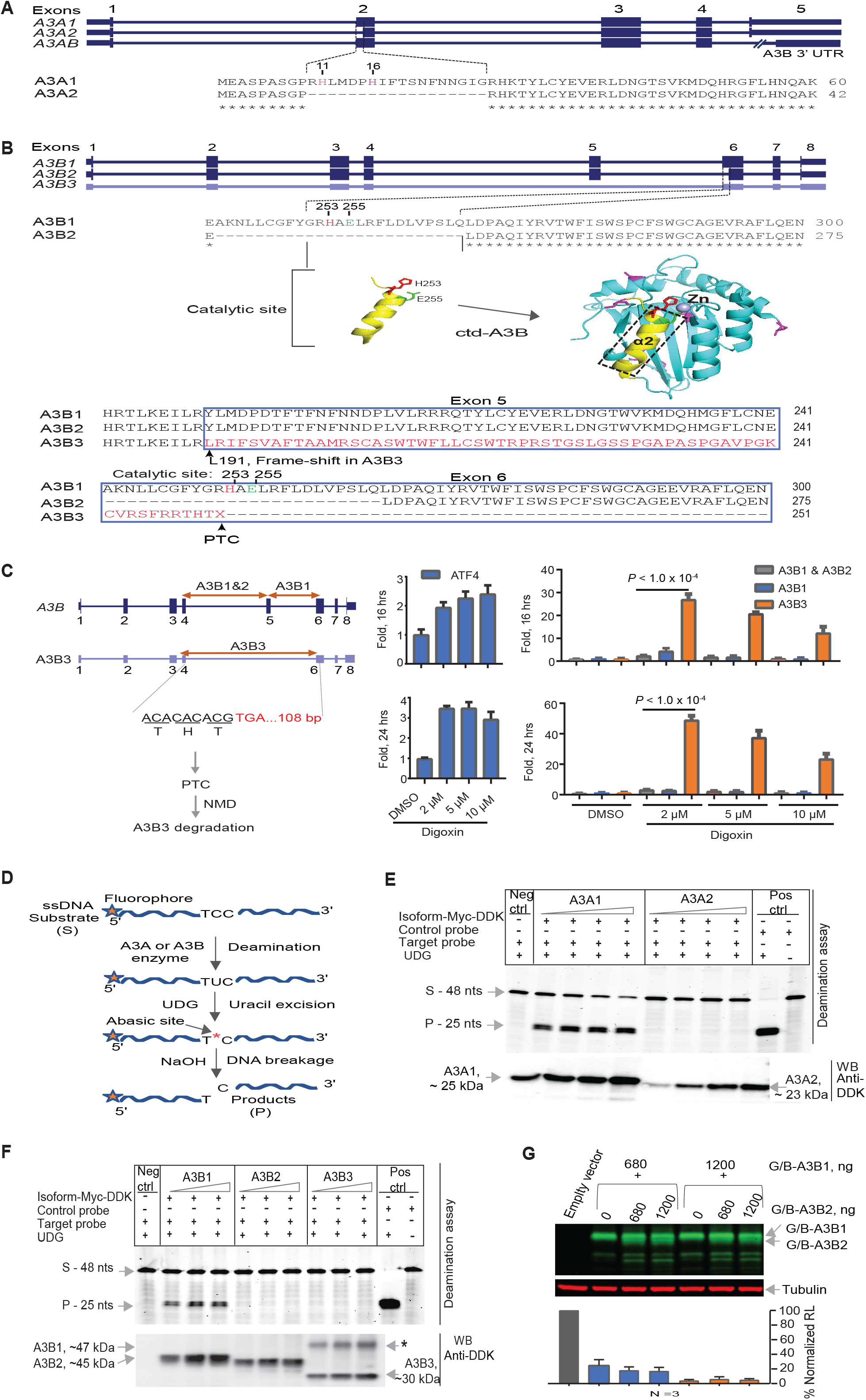
Alternative splicing in *A3A* and *A3B* results in catalytically inactive protein isoforms. Clustal Omega alignment of protein sequences for **A)** canonical A3A1 and alternative A3A2 and **B)** canonical A3B1 and alternative A3B2 and A3B3 isoforms. A3A2 lacks 18 N-terminal aa (R10 - G27), including functional residues H11 and H16^19^, and A3B2 lacks a fragment with functional residues E255 and H253^20^. Skipping of exon 5 generates an *A3B3* transcript that encodes a truncated protein without the catalytic domain, in which a frameshift at position L191 results in the replacement of a 132 aa fragment in the C-terminus of A3B by 62 aa of an aberrant frame-shifted sequence. **C)** Due to a premature termination codon (PTC) in the penultimate exon 6, *A3B3* might be targeted by nonsense-mediated decay (NMD). The effect of NMD was tested by analysis of expression of *A3B* isoforms with TaqMan expression assays indicated by arrows and *ATF4* (positive control) in HT-1376 cells treated with DMSO (vehicle) or digoxin, an NMD-inhibitor. **D)** Outline of the *in vitro* deamination assays testing conversion of ssDNA substrate (S) of 48 nucleotides (nt) into a product (P) of 25 nt by the recombinant C-terminally Myc-DDK tagged **E)** A3A and **F)** A3B protein isoforms. Negative control reactions (Neg ctrl) lack A3A and A3B proteins; positive control probe (Pos ctrl) is completely converted by the UDG enzyme. Western blot (WB) analysis with an anti-DDK antibody shows the amounts of A3A and A3B proteins in reactions. **G**) HIV-1 infectivity restriction assays show no significant inhibitory effects of A3B2 on the activity of the A3B1 protein isoform measured for A3G/A3B protein fusions (G/B, **Figure S3**). Shown one of three representative experiments with Western blot for corresponding recombinant proteins and normalized relative luciferase units (RLU, %) for each labeled condition.

The *A3B3* isoform is generated by skipping of exon 5 in *A3B* and encodes truncated and likely unstable protein without the catalytic domain, in which a 132 aa fragment in the C-terminus of A3B is replaced by 62 aa of an aberrant frameshifted sequence (**Figure 2B**). The stop codon in the penultimate exon also makes the *A3B3* transcript a potential target for NMD. Indeed, expression of *A3B3* mRNA was significantly increased after the treatment of HT-1376 cells with digoxin, a known NMD inhibitor^21^ (**Figure 2C**). Due to NMD, only residual levels of *A3B3* may be detected by expression studies, including by RNA-seq, likely resulting in an underestimation of its expression. Evaluation of the deaminating (mutagenic) potential of recombinant APOBEC3 proteins (**Figure 2D**) showed activities for the canonical A3A1 and A3B1 isoforms, but not for the alternative A3A2, A3B2 and A3B3 proteins (**Figure 2E, F**). The alternative protein isoforms did not show dominant-negative effects on the activities of A3A1 and A3B1 evaluated either by deamination assays (**Figure S2)** or by HIV-1 infectivity inhibition assays^22-24^ (**Figure 2G, Figure S3**). Based on these results, we concluded that AS of *A3A* and *A3B* results in the production of catalytically inactive, non-mutagenic protein isoforms of these enzymes.

### Isoform-specific analysis refines the correlations between *A3A* and *A3B* expression and APOBEC-mediated mutagenesis

Previously, multiple reports showed significant correlations between total expression of *A3A* and *A3B* genes and the burden of APOBEC-signature mutations in several tumor types^8,25^. Now, we revisited these conclusions based on correlations between the expression of the mutagenic *A3A1* and *A3B1* isoforms quantified by RNA-seq read counts for specific exon-exon junctions and ‘APOBEC mutation pattern’ proposed as the most stringent estimate of APOBEC-signature mutation burden^7^. The analysis was performed in six cancer types with ≥10% tumors carrying these mutations (**Figure S4**).

We observed significant positive correlations between the expression of both mutagenic isoforms - *A3A1* and *A3B1* and APOBEC mutation burden in multiple cancers. For *A3A1*, we observed correlations in BRCA (*P* = 5.49E-08; *rho* = 0.17), CESC (*P* = 7.84E-08; *rho* = 0.37) and HNSC (*P* = 3.684E-08; *rho* = 0.24), while for *A3B1 -* in BLCA (*P* = 2.76E-07; *rho* = 0.25). In LUAD, the correlations were comparable for both *A3A1* (*P* = 3.12E-12; *rho* = 0.31) and *A3B1* (*P* = 7.73E-12; *rho* = 0.30) and in LUSC – it was moderately significant only for *A3A1* (*P* = 1.86E-02; *rho* = 0.17) (**Figure 3A, B**). Notably, these tumor-specific correlations were not reported by studies based on total gene expression^8,25^. Our results also corroborate previous findings based on germline variants, which implicated A3A as a more prominent mutagen in BRCA and A3B – in BLCA^1,11^.

**Figure 3:**
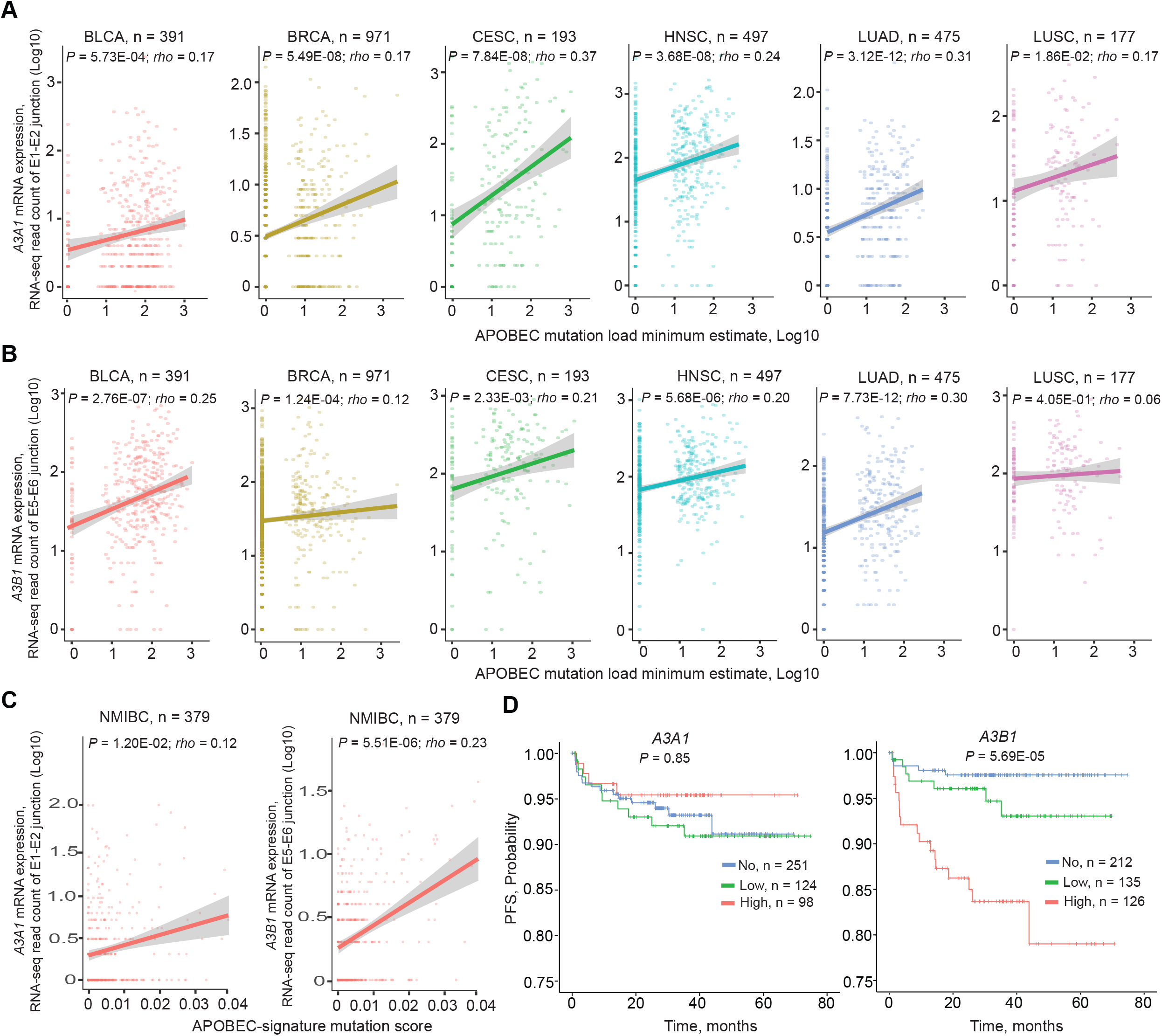
Analysis of *A3A1* and *A3B1* expression based on exon-exon junction RNA-seq reads in relation to APOBEC-mediated mutagenesis and progression-free survival in patients with non-muscle invasive bladder cancer (NMIBC). Correlation analysis between mRNA expression of *A3A1* (**A**) and *A3B1* (**B**) and APOBEC-mediated mutagenesis, measured as ‘APOBEC mutation load minimum estimate’ in six cancer types in TCGA; **C**) in non-muscle invasive bladder cancer (NMIBC) in the UROMOL study, APOBEC-signature mutation score was significantly correlated with *A3B1* but not with *A3A1* expression. **D**) Kaplan-Meier plots for progression-free survival (PFS) of NMIBC based on *A3A1* and *A3B1* isoform expression in the UROMOL study. *P*-values are for multivariable Cox-regression models adjusted for sex, age, and tumor stage. Grouping into “No”, “Low” and “High” groups was done based on *A3B1* RNA-seq read counts, separating samples with no expression (zero reads) and then below and above the median for the remaining samples.

### APOBEC mutagenesis in bladder tumors: A3B as a driver and a predictor of progression-free survival

It was reported that cell lines exhibit episodes of APOBEC-mediated mutagenesis during propagation in culture^26^. The studied cell lines included bladder cancer cell lines, in which we detected the expression of *A3B1* but not *A3A1* (**Note S1**), thus nominating A3B1 rather than A3A1 as the primary mutagenic APOBEC3 enzyme responsible for these episodes of mutagenic activity. Our analysis of non-muscle invasive bladder tumors from the UROMOL study^27^ further supported the role of A3B1 in bladder cancer. Specifically, using the same RNA-seq exon junction isoform-specific quantification of expression as in TCGA samples, we observed that higher expression of *A3B1* was significantly associated with increased APOBEC-mediated mutation burden (*P* = 5.51E-06) and with shorter progression-free survival (*P* = 5.69E-05) in patients with non-muscle-invasive bladder cancer (**Figure 3C, D**). Similar quantification of the *A3A1* expression in the UROMOL tumors showed no association with APOBEC-mediated mutagenesis or progression-free survival.

### The ratio of the non-mutagenic isoform *A3B3* is higher in normal tissues compared to tumors

Alternative *A3B* isoforms (*A3B2* and *A3B3*) are detected in ≥ 50% of 11,058 TCGA samples (**Figure 1**). Although these isoforms may not produce any functional proteins (**Figure 1**), they still could be important. For example, splicing towards alternative, non-mutagenic isoforms might decrease the production of the canonical, mutagenic isoforms. Because mutagenesis is considered tumorigenic, we hypothesized that the proportion of splicing towards non-mutagenic isoforms might be higher in normal tissues compared to tumors. To test this hypothesis, we calculated the percent spliced-in index (PSI)^28^ based on RNA-seq read counts of specific exon junctions in TCGA paired tumor and normal tissues. *A3A* was excluded from this analysis because the expression of the alternative *A3A2* isoform was detected only in 2.57% (18 of 698) of normal samples (**Figure S5A**). Thus, we calculated PSI for exons 5 and 6 of *A3B* in 17 TCGA cancer types with ≥5 tumor/normal pairs. We observed that the proportion of *A3B* alternative splicing was significantly lower in tumors compared to paired normal tissues. Specifically, the proportion of *A3B2* splicing was lower in tumors of KICH (*P* = 1.0E-03) and LUSC (*P* = 3.30E-02) (**Figure 4A, left plot**) and the proportion of *A3B3* splicing was lower in tumors of BLCA (*P* = 4.60E-03), HNSC (*P* = 6.50E-03), LICH (*P* = 1.90E-02), LUAD (*P* = 1.50E-03), and LUSC (*P* = 3.89E-05) (**Figure 4A, right plot**). All other cancer types showed no significant differences in this analysis (**Figure S5B, C**). Most cancer types with a low proportion of non-mutagenic isoforms, specifically of *A3B3*, also had a higher rate of APOBEC-mediated mutation burden^7,8^, with exceptions for tumors of KICH and LICH, in which APOBEC-signature mutations were negligible (**Figure S4)**. In these cancers, A3B might play a mitogenic rather than a mutagenic role as has been shown for hepatocellular carcinoma^29^ and suggested for breast cancer^30^.

**Figure 4.**
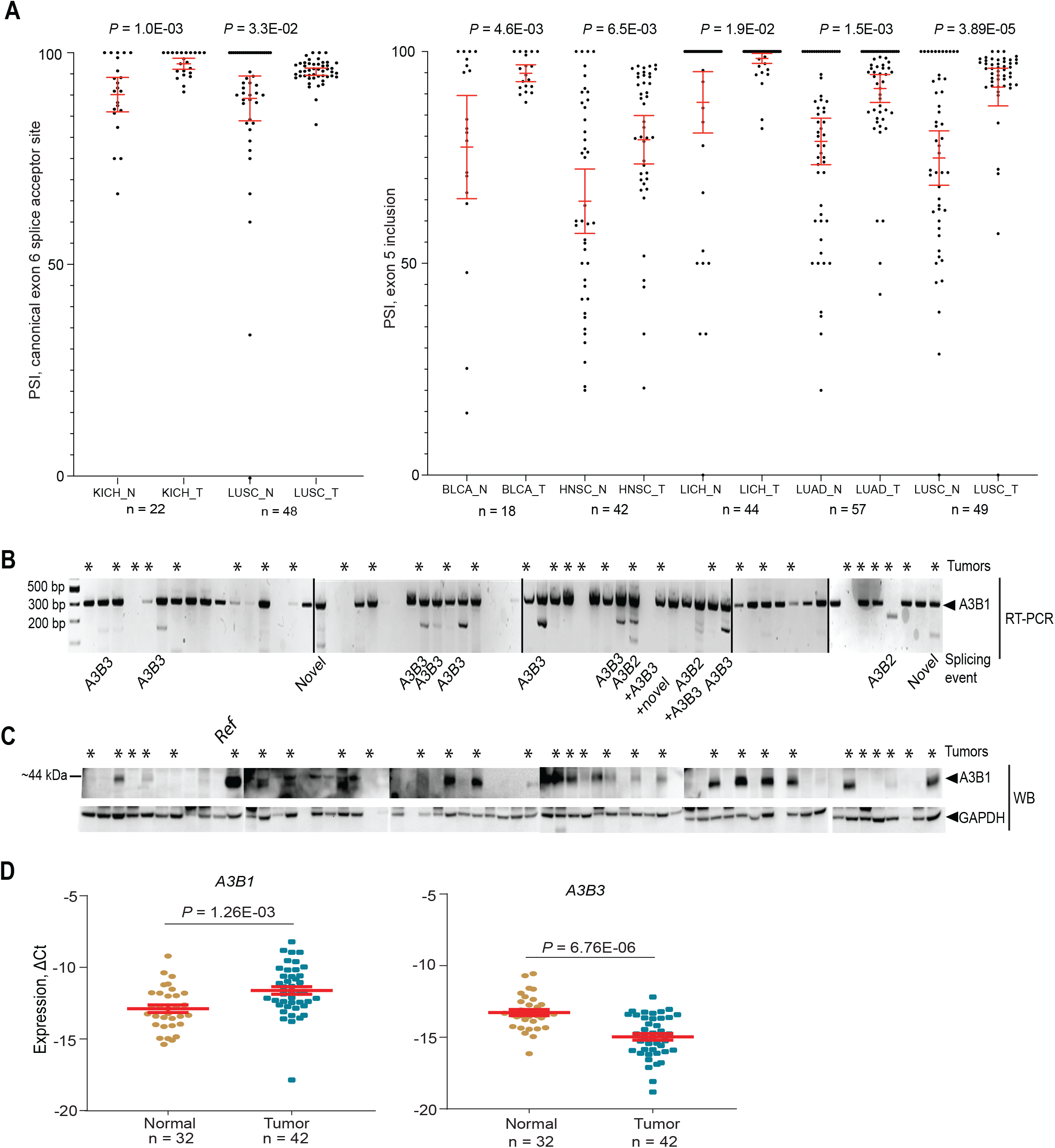
Analysis of *A3B* alternative splicing in paired tumor and adjacent normal samples. The ratios of splicing events generating the canonical mutagenic *A3B1* isoform **-** exon 5 inclusion and the use of canonical exon 6 acceptor site were calculated as percent spliced-in index (PSI) based on RNA-seq reads in TCGA. **A)** Left plot shows that PSI values for the usage of canonical exon 6 acceptor site are significantly higher in tumors compared to normal samples for KICH and LUSC. Right plot shows that PSI values for exon 5 inclusion are significantly higher in tumors compared to normal samples for BLCA, HNSC, LICH, LUAD and LUSC. Red bars represent mean expression levels, *P*-values are for the non-parametric Wilcoxon matched samples signed-rank test. **B**) RT-PCR analysis in 63 bladder tissue samples. Sanger sequencing of RT-PCR products generated with primers for *A3B* exons 4 and 6 shows the canonical (*A3B1*) and alternative (*A3B2* and *A3B3*) splicing events. A splicing event (labeled as “Novel”) that involves both exon 5 skipping and the use of a cryptic splice site in exon 6, was detected in three samples. An AS event (mainly *A3B3*) was observed in 19% of all samples (13 of 63 samples), including 13% of tumors (4 of 30) and 27% of adjacent normal tissues (9 of 33). **C)** Western blot analysis for the A3B1 protein. **D)** qRT-PCR analysis for *A3B1* and *A3B3* isoforms. Mean mRNA expression levels of *A3B1* were significantly higher in tumors and of *A3B3* - in adjacent normal tissues. Red bars represent mean expression levels, *P*-values are for the Student’s t-test.

We also performed qRT-PCR analysis and sequencing of splicing junctions between exons 4 and 6 of *A3B* in a panel of 33 muscle-invasive bladder tumors and 30 adjacent normal tissues (**Figure 4B**). Exon 5 skipping, generating *A3B3*, was the most frequently observed AS event, similar to the pattern observed in TCGA, and more common in normal tissues compared to tumors (**Figure 4B**). Western blot analysis in the same tissue samples showed that A3B1 protein expression was common in tumors but rare in normal tissues, although the frameshifted and truncated A3B3 protein could not be detected (**Figure 4C, Note S2**). We then used isoform-specific TaqMan assays to quantify the expression of *A3B* isoforms in the same set of bladder tissues. The expression of *A3B1* was significantly higher in tumors, while *A3B3* was higher in normal tissues, and *A3B2* was not quantifiable (**Figure 4D**). The fact that AS of *A3B* was more common in normal tissues suggests that increased generation of alternative, non-mutagenic *A3B* isoforms might be anti-tumorigenic and inhibited in tumors.

### *A3B* exon 5 skipping is sensitive to expression levels of some splicing factors

Considering that A3B1 is clinically relevant, at least in bladder tumors (**Figure 3C,D**), decreasing its expression might be of therapeutic importance. We hypothesized this could be achieved by shifting *A3B* pre-mRNA splicing from the mutagenic *A3B1* to non-mutagenic *A3B2* or *A3B3* isoforms. To this end, we first sought to explore the regulation of these splicing events. AS outcomes are regulated by the interaction of *trans*-acting spliceosomal and splicing factors (SFs) with *cis*-acting intronic and exonic pre-mRNA motifs^31^. To explore AS of *A3B*, we created mini-genes by cloning the corresponding alternative exons with 80 bp of flanking intronic sequences into an Exontrap vector (**Figure S6A**). These mini-genes were transiently transfected into HEK293T cells, and their splicing patterns were evaluated. This experimental system was not informative for evaluation of splicing occurring via a cryptic splicing site in exon 6 of *A3B*, as the observed splicing pattern represented only the canonical (*A3B1*) but not the alternative (*A3B2*) isoform (**Figure S6B**). However, we could capture an AS pattern of *A3B* exon 5, with its inclusion representing a proxy of the *A3B1* isoform and its skipping representing a proxy of the *A3B3* isoform. We used this mini-gene construct for exon 5 (E5) to further explore the regulation of *A3B1* versus *A3B3* splicing patterns.

We hypothesized that splicing of *A3B* exon 5 might be affected by SFs that bind within this exon and bioinformatically predicted several candidate SFs (**Table S3**). To experimentally test these predictions, we co-transfected HEK293T cells with expression constructs for 10 of these SFs (**Table S4**) together with the E5 mini-gene; four SFs – SRSF2, SRSF3, CELF1-T4, and ELAVL2 - significantly affected exon 5 splicing (**Figure 5A**), with SRSF2 showing the strongest effect. The effect of SRSF2 on exon 5 splicing was further confirmed in three additional bladder cancer cell lines - HT-1376, SW780, and HTB-9 (**Figure S7**). Screening of exon 5 splicing in 10 cell lines of different tissue origin also showed variable exon 5 skipping (**Figure 5B**), presumably due to differences in levels of expression/activity of endogenous SFs in these cell lines. These results suggested sensitivity of *A3B* exon 5 skipping to expression levels of some SFs, such as SRSF2, which can bind to cis-regulatory motifs^31-34^ within this exon or its flanking introns.

**Figure 5.**
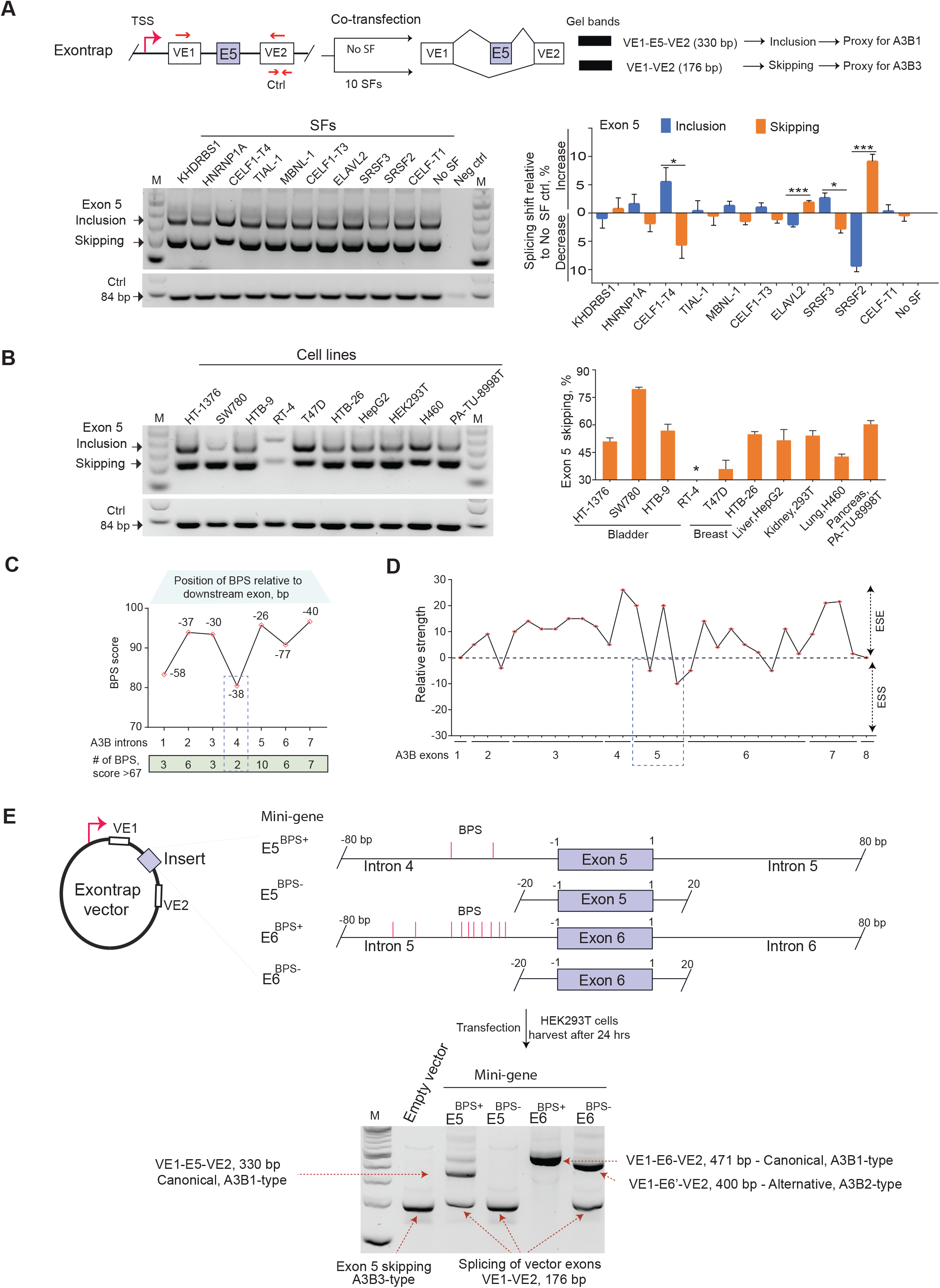
The efficiency of exon 5 skipping depends on branch point sites in intron 4 of *A3B*. **A**) Exontrap mini-gene E5 as an experimental model for the analysis of AS of *A3B* exon 5 (top panel). An agarose gel showing variable ratios of RT-PCR for splicing products of the E5 mini-gene at baseline (No SF control) and after co-transfection with 10 candidate SFs in HEK293T cells; negative control - untransfected cells; M - 100-bp size marker; positive control – vector-specific amplicon of 84 bp. Bands from separate gels representing three biological replicates were quantified by densitometry, and the *A3B1*/*A3B3* splicing shifts were calculated in relation to No SF control (right panel). Overexpression of SRSF2 significantly shifted splicing towards increased *A3B3* expression (about 10%), \**P*<0.05 to \*\*\**P*<0.001 – the significance of *P*-values. **B**) Splicing patterns of the E5 mini-gene analyzed by RT-PCR and agarose gel electrophoresis in 10 human cell lines. RT-4 cell line was excluded due to unspecific PCR products. Bands from separate gels representing three biological replicates were quantified by densitometry to assess the skipping of *A3B* exon 5 (right panel). **C)** Branch points site (BPS) prediction for all 7 introns of *A3B* (**Table S5**). Shown are the BPS with the highest predicted scores for each intron, with indicated positions upstream of the corresponding exons and the numbers of predicted BPS in a 100 bp window. The two predicted BPS within intron 4 have the lowest scores of all *A3B* introns. **D)** Exon splicing enhancer (ESE) and exon splicing silencer (ESS) prediction within *A3B* exons. Exon 5 has the strongest ESS motif of all *A3B* exons. **E**) Comparison of splicing patterns for *A3B* exons 5 and 6 using mini-genes with (E5^BPS+^ and E6^BPS+^) and without (E5^BPS-^ and E6^BPS-^) predicted BPS.

### Skipping of *A3B* exon 5 is facilitated by weak intronic branch point sites within intron 4

Bioinformatics analysis of *A3B* showed differences in the distribution of predicted cis-regulatory motifs - intronic branch point sites (BPS) (**Figure 5C, Table S5**) and exonic splicing silencers/enhancers (**Figure 5D**). The most striking differences were found within intron 4 that harbored the fewest and the weakest scored BPS of all introns in *A3B*. The highest scored intronic BPS located 38 bp and 50 bp upstream of exon 5 were included in the 80 bp of flanking intronic sequences in the E5 mini-gene (referred as E5^BPS+^) and supported the observed partial inclusion of exon 5 that generates the *A3B1*-type canonical splicing event (**Figure 5E**). We then created an E5 mini-gene version with only 20 bp of flanking intronic sequences to exclude the putative BPS and observed no canonical *A3B1*-type splicing in this model (E5^BPS-^, **Figure 5E**), confirming the importance of the putative BPS in intron 4 for exon 5 inclusion/skipping. In agreement with the stronger predicted BPS in intron 5 compared to those in intron 4, only canonical splicing with complete inclusion of exon 6 was observed in a corresponding mini-gene for exon 6 of *A3B* with 80 bp of flanking intronic sequences (E6^BPS+^). Consistently, no canonical splicing was observed in the E6^BPS-^ mini-gene, which lacks the intron 5 BPS (**Figure 5E**).

Based on these results, we concluded that skipping of *A3B* exon 5 is facilitated by weak BPS in intron 4. The magnitude of exon 5 skipping is likely to be determined by expression levels of some SFs that might differ between tissue types and disease conditions, including normal tissues vs. tumors.

### SF3B1 inhibitor pladienolide B promotes skipping of *A3B* exon 5 and reduces A3B1 production

Efficient canonical splicing requires robust interaction of the spliceosomal machinery with intronic BPS; thus, exon 5 skipping could represent a weak engagement of the spliceosome to BPS in intron 4 (**Figure 6A**). Spliceosomal interaction with BPS is facilitated by the SF3b complex, with SF3B1 as a core protein^35^. We hypothesized that an inhibitor of SF3B1, such as pladienolide B^35,36^, would destabilize interaction of SF3b complex at weak BPS within intron 4, resulting in enhanced exon 5 skipping, but may not significantly affect exon 6 splicing that is regulated by strong BPS in intron 5 (**Figure 6B**). We tested this hypothesis by evaluating splicing patterns of E5^BPS+^ and E6^BPS+^ mini-genes in cells treated with pladienolide B. This treatment not only enhanced but caused complete exon 5 skipping in a concentration-dependent manner while not affecting exon 6 splicing (**Figure 6C**). Skipping of endogenous exon 5 of *A3B* was also increased by this treatment, resulting in higher expression of *A3B3* mRNA and reduced expression of *A3B1* mRNA and A3B1 protein (**Figure 6D, E, Figure S8A**). Importantly, the reduction of A3B1 protein caused by pladienolide B treatment was more prominent than the induction of *A3B3* expression, although it might be difficult to detect due to degradation of *A3B3* transcript by NMD (**Figure 1C)**. This hypothesis was supported by a much stronger induction of *A3B3* expression by pladienolide B in the presence of digoxin, an NMD blocker^21^ (**Figure 6F**). These experiments confirmed that exon 5 skipping is facilitated by weak BPS in intron 4 but is dependent on the availability of active SFs, such as SF3B1. The role of SF3B1 in this process was further confirmed by its siRNA-mediated knockdown in HT-1376 cells, resulting in increased skipping of *A3B* exon 5 (**Figure S8B**). Thus, sequestering of active SF3B1 by any of its inhibitors should increase *A3B* exon 5 skipping, leading to reduced levels of mutagenic A3B1.

**Figure 6.**
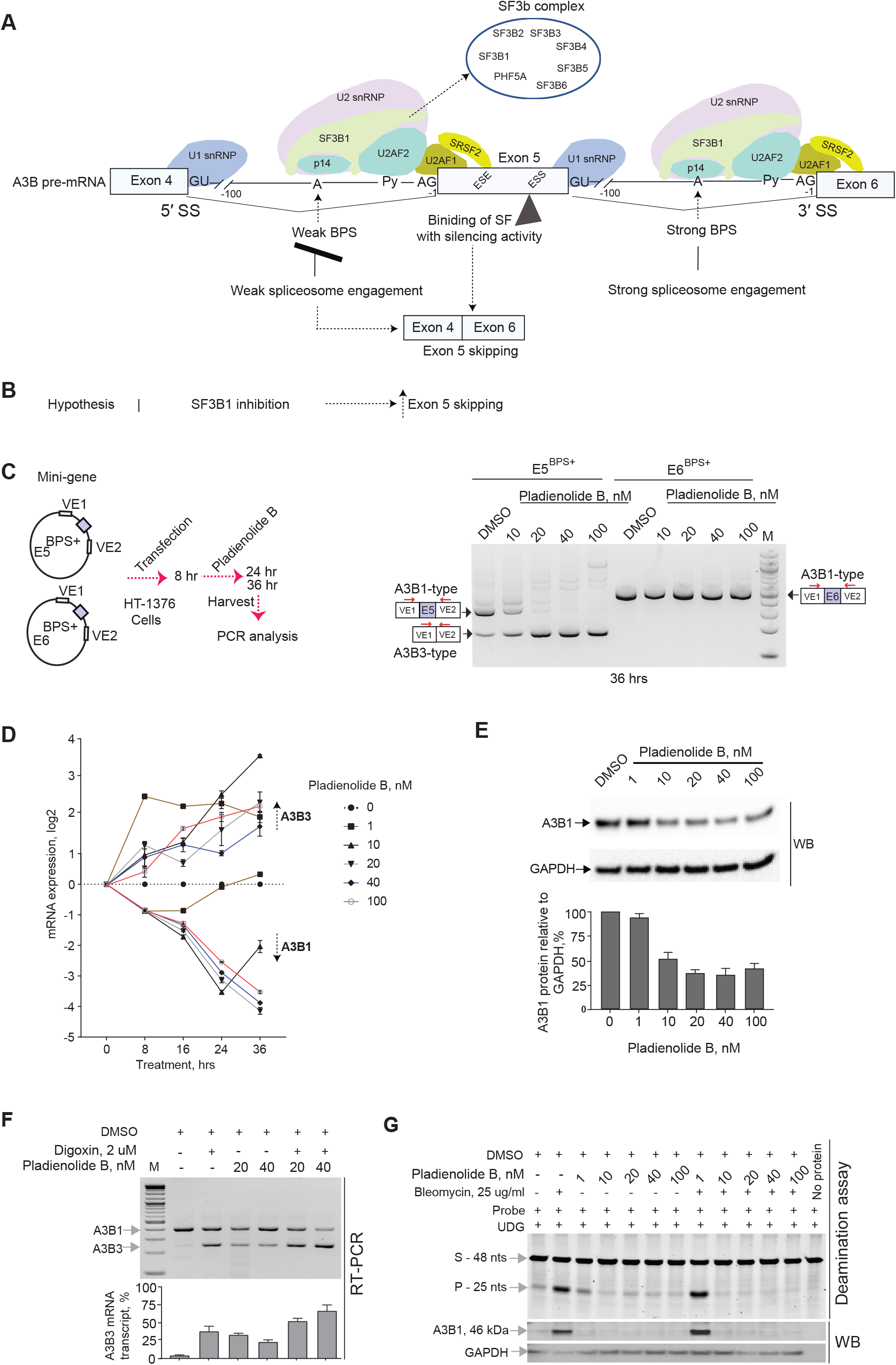
Modulation of *A3B* exon 5 skipping by an SF3B1 inhibitor pladienolide B. **A**) Schematic representation of the *A3B* splicing module (exon 4 - exon 6) featuring interactions between *trans*-factors of the spliceosomal machinery with splicing *cis*-elements: splice sites (SS), branch point sites (BPS), polypyrimidine tract (Py), and exonic splicing enhancers/silencers (ESE)/(ESS); possible outcomes of *A3B* exon 5 splicing resulting from interactions between SF3B1 and other spliceosomal proteins with weak versus strong BPS. **B**) Schematic representation of the hypothesis depicting that inhibition of SF3B1 is expected to result in skipping of *A3B* exon 5 and downregulation of A3B1 protein. **C**) PCR analysis of splicing patterns of *A3B* exons 5 and 6 using Exontrap mini-genes E5^BPS+^ and E6^BPS+^ in HT-1376 cells treated for 36 hrs with increasing concentrations of pladienolide B. Exon 5 skipping is increased in the presence of the SF3B1 inhibitor, while splicing of exon 6 is not affected. **D**) Results of qRT-PCR in HT-1376 cells treated with increasing concentrations of pladienolide B for indicated time points show splicing shift from *A3B1* to *A3B3*. **E)** Results of RT-PCR expression analysis and Western blot protein analysis of HT-1376 cells treated with increasing concentrations of pladienolide B for 36 hrs show splicing shift from *A3B1* to *A3B3*, resulting in downregulation of A3B1 protein, while the aberrant and unstable A3B3 protein was not detected. Protein quantification is for three independent experiments. **F)** RT-PCR analysis of exon 5 splicing representing *A3B1* and *A3B3* transcripts in HT-1376 cells treated with DMSO (vehicle), digoxin (an NMD-inhibitor), pladienolide B alone, and pladienolide B with digoxin. *A3B3* mRNA levels are increased in digoxin-treated cells due to its inhibition of NMD. **G**) Deamination assays using whole-cell extracts from HT-1376 cells treated with bleomycin and pladienolide B. Equal amounts of total protein were used for each reaction based on densitometry pre-assessment of GAPDH protein levels in each sample (not shown). Deamination was observed in reactions with protein lysate of HT-1376 cells treated with bleomycin alone or bleomycin plus 1 nM of pladienolide B. Higher concentrations of pladienolide B (10, 20, 40 and 100 nM) significantly reduced A3B1 levels and prevented deamination of ssDNA probe in bleomycin-treated conditions. Western blot (WB) analysis with an anti-A3B1 antibody shows the amounts of A3B1 in corresponding reactions. A3B1 was barely detectable in DMSO and pladienolide B only conditions because of very low total protein input in reactions (2 ul of total lysate). Shown are representative results of one of the three independent experiments.

Reducing the levels of mutagenic A3B1 enzyme might restrict APOBEC mutagenesis in some clinically relevant conditions. To test this, we simultaneously treated cells with bleomycin, a chemotherapy drug, which induces *A3B* expression^1^, and pladienolide B. Deaminating (mutagenic) activity observed in lysates of these cells due to the presence of endogenous bleomycin-induced A3B was completely blocked by pladienolide B (**Figure 6G**). We also showed that pladienolide B inhibited APOBEC-mediated mutagenic activity in a cell-based cytosine deamination assay (**Figure S9**).

## DISCUSSION

High activity of the A3A and A3B enzymes is mutagenic and genotoxic, as suggested by *in vitro* overexpression studies^38-41^ and significant positive correlations between mRNA expression of genes encoding these enzymes and burden of APOBEC-signature mutations in tumors^1,8,25,42^. Multiple mechanisms likely exist to regulate the activity of these enzymes and restrict cell damage they may cause. Here, we showed that AS of *APOBEC3s*, particularly of *A3B*, is one of several possible regulatory mechanisms controlling the expression of the mutagenic A3B1 enzyme. We present proof-of-principle data suggesting that AS can be modulated to shift the A3B balance from producing mutagenic to non-mutagenic isoforms.

Initially, we explored the functional consequences of splicing events within exon 2 in *A3A* and exon 6 in *A3B*, both occurring via cryptic splicing sites, and skipping of the entire exon 5 in *A3B*. In all these cases, AS resulted in a shift from the production of the canonical isoforms (*A3A1* and *A3B1*) that encode mutagenic enzymes, to corresponding alternative isoforms (*A3A1, A3B2*, and *A3B3*) that encode non-mutagenic enzymes. The expression of the canonical isoforms positively correlated with a load of APOBEC-signature mutations in TCGA tumors of different types. The isoform-level analysis also suggested that *A3A1* and *A3B1* might contribute to APOBEC mutagenesis in a cancer-type specific manner. Our previous finding that a common genetic variant identified by a GWAS for bladder cancer risk was also associated with *A3B* expression and APOBEC mutagenesis^1^, nominated A3B as the primary mutagenic APOBEC in bladder tumors. This was further supported by the association of increased *A3B1* expression with higher APOBEC mutation score and shorter progression-free survival in patients with non-muscle invasive bladder cancer.

AS is regulated by a complex interplay between the core spliceosomal and alternative SFs^31^ that bind cis-acting exonic and intronic elements. These interactions can depend on various tissue- and disease-specific environments^43^. Analysis of splicing patterns using our *in vitro* mini-gene Exontrap system identified splicing plasticity of *A3B* exon 5, which we attributed to the weak BPS in intron 4 of this gene and levels of SF3B1. Splicing of endogenous exon 5 also varied in cell lines of different tissue origin and was sensitive to changes in expression levels of some other SFs predicted to bind within this exon, such as SRSF2.

We observed that *A3B* splicing events were more common in adjacent normal tissues compared to tumors of several types. This suggests that AS of *A3B* is an intrinsic, tissue-specific regulatory mechanism rather than a result of general dysregulation of splicing machinery in tumors^44,45^, manifested in the inactivation of tumor suppressor genes and generation of tumor-specific isoforms^46^. On the other hand, mutations in splicing or other regulatory factors that would affect splicing globally, could also result in decreased expression of alternative *A3B2* and *A3B3* isoforms and, consequently, lead to increased APOBEC mutagenesis. Thus, AS of *A3B* might be a natural mechanism restricting the expression of mutagenic isoforms in some conditions such as in normal tissues.

The observed splicing plasticity of *A3B* mRNA might represent an adaptive biological mechanism of tuning down the excessive effects of mutagenic APOBEC3 proteins to safeguard the cells from the genotoxic activity of these enzymes. A similar role has been proposed for several SFs, including SRSF2, which affected *A3B* exon 5 splicing in our experiments. These SFs regulate the expression of DNA repair proteins to protect the genome from DNA damage and the toxic effects of mutagens^47-50^. Splicing re-routing, such as by exclusion of exon 5 in *A3B*, followed by elimination of the alternative frame-shifted *A3B3* transcript by NMD might be a mechanism to tweak APOBEC mutagenesis in specific conditions. The entire functional role of *A3B3* could be just to use up some pre-mRNA that otherwise would be used to produce the mutagenic A3B1 enzyme, and then get degraded by NMD. Thus, the NMD-targeted *A3B3* transcript not producing a functional protein might still play an important role in the regulation of APOBEC mutagenesis, regardless of its low residual expression levels observed in TCGA tumors.

A recent study has analyzed mutational signatures in a large set of cell lines and suggested that the initiators of APOBEC mutagenesis *in vitro* are cell-intrinsic factors with continuous but intermittent activity^26^. As the authors did not observe significant correlations between APOBEC mutagenesis and expression of *APOBEC3* genes, they suggested that these initiators may include modulators such as the availability of single-stranded DNA (ssDNA) substrate, etc. Based on our observations that *A3B1* and not *A3A1* is expressed in bladder cancer cell lines, which have a high load of APOBEC-signature mutations, it is likely that at least in these cell lines episodic APOBEC-mediated mutagenesis is caused by the A3B1 activity.

Therapeutic targeting of AS to treat cancer is a rapidly developing field^51^ with several types of SF3B1 inhibitors, such as E7107^52,53^, an analog of pladienolide B, and H3B-8800^54^ being evaluated for acute myeloid leukemia (AML) and myelodysplastic syndrome (MDS)^36,51,54^. Because tumor cells depend on wild-type SFs for survival, these drugs are particularly effective in killing cancer cells with already impaired splicing machinery due to mutations in SFs, such as SRSF2 and SF3B1^54^. Mutations in these and other SFs have been reported in many solid tumors, including bladder cancer^55^ and these tumors tend to have more APOBEC-signature mutations (**Figure S10**). Our results suggest that SF3B1 inhibitors and other tools affecting alternative splicing, might be tested to eliminate cancer cells with mutations in SF and also to control APOBEC mutagenesis in clinically relevant conditions. It is interesting to note that *FGFR3*-S249C, the most common somatic mutation found in the highly recurrent non-muscle invasive bladder cancer, is likely caused by APOBEC mutagenic activity^56^. As we found A3B1 the primary mutagenic APOBEC enzyme in bladder tumors, enhancing *A3B* exon 5 skipping might help to restrict APOBEC-mediated mutagenesis, including *FGFR3*-S249C mutation in bladder cancer, as well as prevent tumor progression and recurrence, clonal evolution, and resistance to chemotherapy (**Figure 7**).

**Figure 7.**
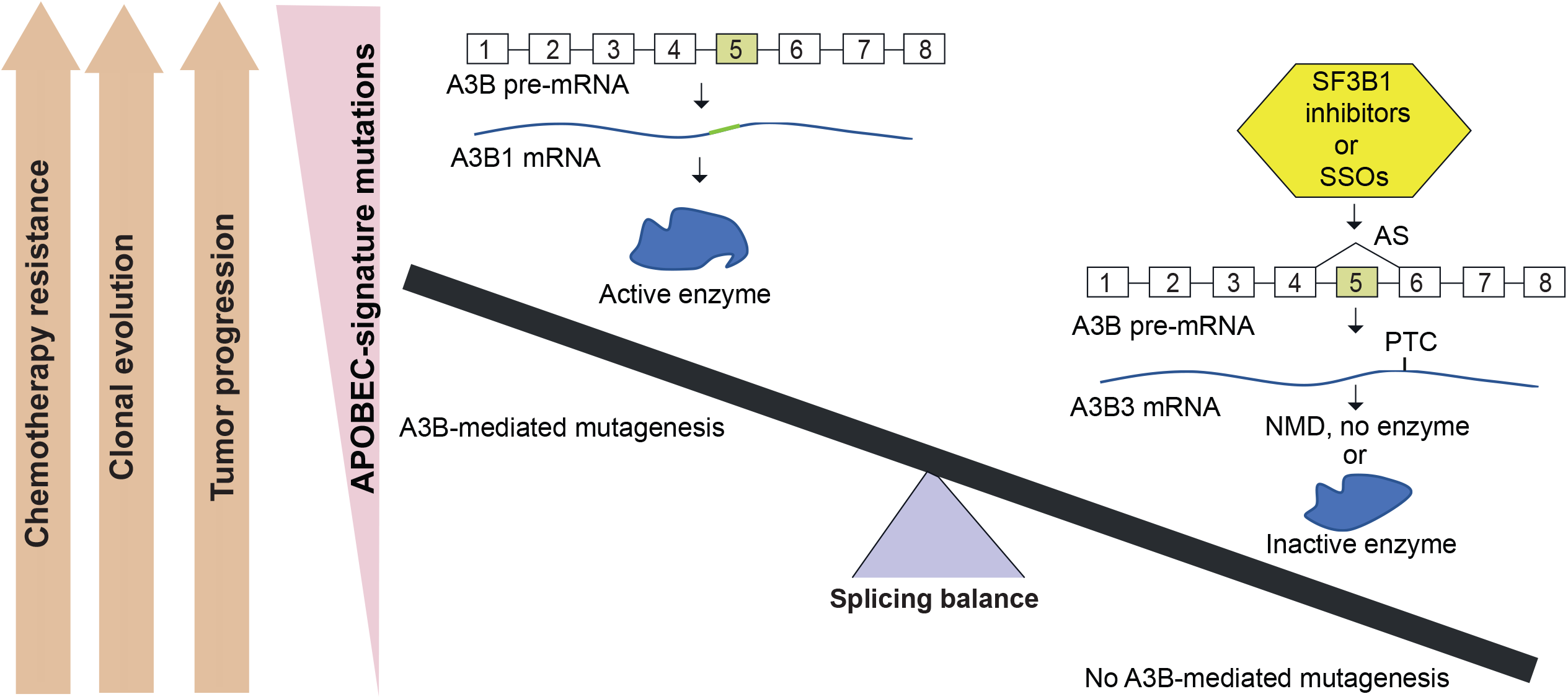
The proposed role of alternative splicing of *A3B* and its targeting for modulating APOBEC mutagenesis. Canonical splicing of *A3B* pre-mRNA generates the mutagenic A3B1 enzyme that causes APOBEC mutagenesis and fuels tumor progression, recurrence, clonal evolution, and chemotherapy resistance. Alternative splicing of *A3B*, and, specifically, skipping of exon 5, generates *A3B3* isoform with a premature termination codon (PTC) that is degraded by nonsense-mediated decay (NMD); the residual transcript encodes catalytically inactive protein isoform. Therapeutic enhancing of exon 5 skipping by SF3B1 inhibitors or other tools affecting alternative splicing such as splicing-switching oligonucleotides (SSOs), may restrict and even prevent A3B-mediated mutagenesis in clinically relevant conditions.

In conclusion, our study showed that AS of *A3B*, which creates non-mutagenic isoforms, represents an intrinsic regulatory mechanism that keeps in check the expression of the mutagenic A3B1 enzyme, modulates APOBEC-mediated mutagenesis in different disease-, tissue- and environment-specific conditions, and can be therapeutically targeted. Additional functional studies will be needed to better understand the consequences of this modulation for clinical outcomes.

## MATERIALS AND METHODS

### Analysis in TCGA samples

#### Data acquisition and processing

The mutation dataset for all TCGA cancer types was downloaded from the Broad GDAC Firehose in October 2016. For analysis of APOBEC-signature mutations, we used the variable “APOBEC mutation load minimum estimate”, which represents APOBEC mutation pattern^8^. mRNA expression of *A3A* and *A3B* genes was analyzed in 11,058 TCGA samples (10,328 tumors and 730 adjacent normal tissues) based on RNA-seq BAM slices generated using workflow https://docs.gdc.cancer.gov/API/Users_Guide/BAM_Slicing/ through the NCI Genomics Data Commons (GDC) portal accessed in August 2019.

#### Estimation of A3A and A3B RNA-seq read counts for exons and exon-exon junctions

For estimation of sequencing reads corresponding to all exons as well as canonical and alternatively spliced exon-exon junctions, RNA-seq Bam slices were processed through R package ASpli version 1.5.1. We used strict ASpli pipeline settings, requiring a minimum of 10 nucleotides of perfect match covering exon-exon junctions.

#### Correlation analysis of mRNA expression and APOBEC mutagenesis

Expression levels of the mutagenic isoforms *A3A1* and *A3B1* were calculated based on their RNA-seq exon-exon junction read counts. The counts of sequencing reads for E1-E2 junctions (for *A3A1*) and E5-E6 junctions (for *A3B1*) and APOBEC mutation load minimum estimate counts were Log10-transformed (after adding 1 to all raw values). The Spearman correlation analysis and data plotting were performed using R packages for six cancer types with ≥10% of samples with APOBEC signature mutations. SKCM was excluded from the analysis because mutations induced by APOBEC in melanoma likely overlap with mutations induced by UV^57^.

#### Analysis of AS levels in paired tumor and adjacent normal tissue samples

The analysis was performed in 17 cancer types with ≥5 tumor-normal pairs. Splicing ratios were calculated as PSI based on RNA-seq reads using ASpli pipeline, as was previously described^28^ and statistical significance between PSI in tumors vs. normal tissues was evaluated by non-parametric Wilcoxon matched pairs signed rank test.

### RNA-sequencing of SeV-infected T47D cells

Total RNA extracted from the T47D breast cancer cells infected or not infected (control) with Sendai Virus (SeV)^1^, was used for paired-end RNA-seq on HiSeq 2500 (Illumina) in biological duplicates. The library was prepared with KAPA Stranded RNA-seq Kit with RiboErase (Roche). RNA-seq reads (120 bp) were filtered and aligned with STAR alignment tool^58^ using the GRChg37/hg19 genome assembly and visualized using the Integrative Genomics Viewer (IGV). The RNA-seq reads aligned to *A3A* were Bam-sliced with SAM tools and then re-aligned to the reference genome by STAR to identify cross-alignment with *A3B*.

### Bioinformatics analysis of the A3A and A3B protein isoforms

Protein sequences of the APOBEC3 isoforms A3A1 (UCSC ID uc003awn.2), A3A2 (UCSC ID uc011aob.1), A3B1 (UCSC ID uc003awo.1), A3B2 (UCSC ID uc003awp.1), and A3B3 (UCSC ID uc003awq.1) were downloaded from the UCSC genome browser. Multi-sequence alignments were generated using the web-based tool Clustal Omega^59^.

### Cell lines

Cell lines - embryonic kidney HEK293T, bladder cancer cell lines - HT-1376, RT-4, HTB-9, and SW780, breast cancer cell lines - MCF-7, MDA-MB-231 (HTB-26), and T47D (HTB-133), hepatocellular carcinoma HepG2, and lung carcinoma H460 - were purchased from the American Type Culture Collection (ATCC) and maintained per ATCC recommendations. A pancreatic cancer cell line PA-TU-8998T was purchased from Leibniz Institute DSMZ-German Collection of Microorganisms and Cell Cultures (DSMZ Scientific). No commonly misidentified cell lines were used in this project. If used longer than for a year after initial purchase, cell lines were authenticated by the Cancer Genomics Research Laboratory of NCI by genotyping a panel of microsatellite markers (Identifiler kit, Thermo Fisher Scientific). All cell lines were tested bi-monthly for mycoplasma contamination using the MycoAlert Mycoplasma Detection kit (Lonza).

### Analysis in non-muscle invasive bladder tumors from the UROMOL study

A set of low-stage (Ta and T1) bladder tumors representing non-muscle-invasive bladder cancer (NMIBC) has been described^27^. Mutations were scored based on RNA-seq data and used for deconvolution into mutational signatures S1-S6, with S3 corresponding to the APOBEC-signature mutations^27^. The FASTQ files for RNA-seq data were aligned with STAR and BAM-sliced to include *A3A* and *A3B* genes, followed by estimation of all exon and exon junction reads using ASpli R package, similar to the analysis in TCGA samples. We performed a Spearman correlation analysis of log10-transformed read counts for *A3A1* and *A3B1* with APOBEC-signature mutation score (S3). Progression-free survival analysis was performed based on the expression of *A3A1* and *A3B1* mutagenic isoforms with samples divided into three groups: “No” - samples with undetectable expression (0 value); the remaining samples were split into two groups based on the expression below and above the median as “Low” and “High” groups, respectively. The Cox-regression models were adjusted for sex,age, and tumor stage.

### Analysis of additional bladder tumor and adjacent normal tissue samples

The panel of muscle-invasive bladder tumors (n = 42) and adjacent normal (n = 32) tissue samples has been described^60^. For each of these samples, cDNA was synthesized from 250 ng of total RNA per 20 µl reactions using the RT^2^ First-Strand cDNA kit and random hexamers (Qiagen). For detection of splicing events between *A3B* exons 4 and 6, we performed PCR with primers: F_ex4: 5’-GCCTTGGTACAAATTCGATGA-3’ and R_ex6: 5’-TGTGTTCTCCTGAAGGAACG-3’, with cDNA input corresponding to 30 ng of total RNA per 25 ul reactions using AmpliTaq Gold™ 360 Master Mix. PCR-amplified products were resolved on 2% agarose gel, and each distinct PCR product was cut, purified, and Sanger-sequenced. For quantification of *A3B1* and *A3B3* splicing products, cDNA input corresponding to 10 ng of total RNA per reaction was used, as previously described^1^, and subjected to qRT-PCR using isoform-specific TaqMan assays (**Note S4**). Expression of *A3B1* and *A3B3* was normalized by the expression of endogenous controls *GAPDH* (assay 4326317E) and *PPIA* (assay 4326316E). Western blot analysis for A3B1 protein and GAPDH (loading control) was performed as described in **Note S2**.

### Generation and partial purification of the recombinant A3A and A3B protein isoforms

Expression constructs for the C-terminally Myc-DDK tagged canonical isoforms *A3A1* (NM_145699) and *A3B1* (NM_004900) cloned in the pCMV6 vector were purchased from OriGene (Rockville, MD). Open reading frames for the C-terminally Myc-DDK tagged alternative isoforms *A3A2, A3B2*, and *A3B3* (**Table S1**) were synthesized (Thermo Fisher Scientific) and cloned into BamHI/XbaI restriction sites of the pcDNA3.1(+) vector. The HEK293T cells (4 x10^6^ cells/20 ml) were seeded in 175cm^2^ flasks (Corning) and transiently transfected with plasmids after 24 hrs at 75% confluency using Lipofectamine 3000 (Thermo Fisher Scientific). Cells were harvested 24 hrs post-transfection and proteins were purified with c-Myc tagged Protein Mild Purification Kit (MBL, Japan) and eluted with 20 μl of 1 mg/ml Myc peptide provided with the kit. The concentration of the total eluted protein, which included both purified protein and Myc peptide, was estimated using a BCA protein assay (Thermo Fisher Scientific). For evaluating protein purity and enrichment, ∼25 ug of total protein was resolved on 4–12% Tris-glycine SDS polyacrylamide gel (Life Technologies) and used for Western blot analysis. Densitometry analysis of Western blots showed at least 10-fold enrichment of all eluted isoforms compared to whole-cell lysates (**Note S3**). All candidate antibodies were tested for the detection of A3A and A3B protein isoforms after overexpression of corresponding expression constructs in cell lines (**Note S2)**. Detection was done using HyGLO chemiluminescent HRP antibody detection reagent (Denville Scientific Inc).

### Cytosine deamination assays with recombinant proteins

Deamination activity of the recombinant A3A and A3B protein isoforms was tested as previously described^61^. Specifically, reactions were carried out in 10 µl of deamination buffer containing 10 mM Tris/HCl, pH 7.5, 50 mM NaCl, 1 mM DTT, 0.25 µg - 1 µg of partially purified A3A and A3B proteins and 1-5 pM of single-stranded DNA substrate – the target probe 5’-5Alexa488N/(ATA)_8_<u>TCC</u> (ATA)_7_-3’, or a positive control probe 5’-5Alexa488N/(ATA)_8_<u>TUU</u> (ATA)_7_-3’ (Invitrogen). Reactions were incubated in a water bath at 37°C for 2 hrs, treated with the Uracil DNA Glycosylase (UDG) for 40 min at 37°C, and then with 0.6 N NaOH for 20 min at 37°C. Final products were mixed with 2x RNA loading dye (Thermo Fisher Scientific) and heated at 95°C for 2-3 min. Of the final 40 ul reaction volume, one set of 14 µl aliquots was resolved on 15% TBE-urea polyacrylamide gel (Life Technologies) at 150 V for 1 hrs and 20 min at room temperature in 1x TBE buffer. Gels were imaged with Gel Doc (Bio-Rad) using Alexa 488 fluorescence settings. Another set of 14 µl aliquots from the same reactions was used for the detection of APOBEC3 proteins by Western blotting. Concentrations of eluted proteins were estimated based on densitometry of Western blots. For competition assays, the amounts of mutagenic isoforms and total reaction volumes were kept constant while the amounts of non-mutagenic isoforms were increased. Proteins extracted from the lysates of untransfected cells were used as a negative control to account for inhibition caused by non-specific endogenous proteins. In the 1:1 control competition reaction, the non-mutagenic isoform was replaced by an equal amount of the negative control protein.

Deamination activity of endogenous A3B1 in the presence of pladienolide B was evaluated using a previously described protocol^14^ with some modifications. Briefly, HT-1376 cells, treated with bleomycin (25 µg/ml) alone, pladienolide B alone (1, 10, 20, 40, 100 nM) and with combined bleomycin and pladienolide B, were harvested after 48 hrs in HED buffer (25 mM HEPES, 5 mM EDTA, 10 % glycerol, 1 mM DTT and protease inhibitor). The protein concentrations were determined based on a densitometric assessment of GAPDH by Western blot from 7 ul of total lysate for each condition. Each 20 ul deamination reaction contained equal amounts of total protein in 15 ul (adjusted with H2O), 1 ul (10 pM) of single-stranded DNA substrate –probe 5’-5Alexa488N/(ATA)_8_<u>TCC</u> (ATA)_7_-3’, 2 ul 10× UDG buffer (Thermo Scientific), 1 ul (1U/ul) UDG and 1 ul RNaseA (100 mg/ml, Qiagen) and reactions were incubated at 37°C for 3 hrs. Subsequently, 100 mM NaOH was added to each reaction, and the samples were then incubated at 37°C for 30 minutes to cleave the abasic sites. Final products were mixed with 2x RNA loading dye (Thermo Fisher Scientific) and heated at 95°C for 2-3 min, and reactions were resolved on a 15 % urea-TBE gel and imaged as described above.

### HIV-1 infectivity inhibition assays

The activity of the A3B protein isoforms was evaluated with single-cycle infection assays for HIV-1 restriction as has been described for APOBEC3G (A3G)^22-24^. Briefly, a G/B-A3B1 plasmid was constructed starting from the previously engineered A3B plasmid^22^ by replacing 63 aa at the N-terminus with a similar region of A3G. This replacement increases the packaging of A3B protein into HIV-1 viral particles, which is necessary for the ability of A3B to inhibit HIV-1 infectivity^22,24^. This system is suitable for analysis of A3B activity, which, like A3G, has two cytidine deaminase domains, but is not appropriate for evaluating the activity of A3A as it contains only one cytidine deaminase domain and A3G/A3A swap would not be compatible. Fusion constructs G/B-A3B-V1, G/B-A3B-V2, G/B-A3B-V3, were generated to represent three A3B expression constructs, all with C-terminal hemagglutinin epitope tags (3X-HA). All plasmids were verified by Sanger sequencing. Because the A3B plasmids were from three sources, we found two protein-changing single point variations that might be functionally relevant (**Figure S3**). To generate the virus for infection, HEK293T cells (4 × 10^5^ cells/ 6-well dish) were transfected using LT1 reagent (Mirus Bio) with HDV-EGFP (1 ug), pHCMV-G (0.25 ug), and variable concentrations of plasmids (0, 680 and 1200 ng), individually or in combinations. The virus was harvested 48 hrs post-infection, filtered with 0.45-um-pore filters, and stored at −80 °C. Capsid p24 measurements were determined using the HIV-1 p24 capsid (CA) ELISA Kit (XpressBio). Normalized p24 CA amounts were used to infect TZM-bl cells containing HIV-1 Tat-inducible luciferase reporter gene, in a 96-well plate (4000 cells/well). Luciferase activity was measured 48 hrs after infection, using a 96-well luminometer (LUMIstar Galaxy, BMG LABTECH). For some experiments, portions of the viral supernatant were spun through a 20% sucrose cushion (15,000 rpm, 2 h, 4°C, in a Sorvall WX80 + ultracentrifuge), concentrated 10-fold, and used in experiments to determine virion encapsidation of APOBEC3 proteins by Western blotting analysis as previously described^23^.

### Evaluation of *A3B3* mRNA degradation by nonsense-mediated decay (NMD)

The HT-1376 cells were treated with DMSO (vehicle) or with 2, 5, and 10 µM of digoxin (Sigma), an NMD inhibitor^21^ dissolved in DMSO. The cells were harvested after 16 and 24 hrs of treatment; total RNA was isolated with an RNeasy kit with on-column DNase I treatment (Qiagen), and RNA quantity and quality were analyzed with NanoDrop 8000 (Thermo Scientific). After an additional DNA removal step, cDNA for each sample was prepared from equal amounts of total RNA, using the RT^2^ First-Strand cDNA kit and random hexamers (Qiagen). Expression was measured in the same cDNA with TaqMan expression assays (all from Thermo Fisher) for endogenous controls *GAPDH* (assay 4326317E) and *PPIA* (assay 4326316E), and positive control *ATF4* (assay Hs00909569_g1), which is induced by NMD inhibition^62^; custom assays were used for detection of *A3B1, A3B3*, and *A3B1/A3B2* combined (**Note S4**). Experiments were performed in biological triplicates per condition and expression was measured in four technical replicates on QuantStudio 7 (Life Technologies) using TaqMan Gene Expression buffer (Life Technologies). Water and genomic DNA were used as negative controls for all assays. Expression was measured as *C*_t_ values (PCR cycle at detection threshold) and calculated as fold change using 2^-(ΔΔ*C*t)^ method in relation to control (untreated) groups of samples.

### Bioinformatics analysis of *A3B* splicing *cis*-elements and SF binding sites

Exonic sequences and 100 bp of intronic sequences upstream of each exon were used for the prediction of exonic splicing enhancer (ESE)/ silencer (ESS) motifs and branch point sites (BPS) using the Human Splicing Finder (HSF, www.umd.be/HSF3/)^63^. Per HSF guidelines, BPS with scores above 67 were considered high-confidence; the strength of ESE and ESS was evaluated based on the relative ESE/ESS ratio. Splicing factor (SF) binding sites were predicted using SFmap^64^ and SpliceAid2^65^ (**Table S3**).

### Exontrap analysis of alternatively spliced exons of *A3A* and *A3B* genes

Exon 2 of *A3A* and exons 5 and 6 of *A3B* with either 20 or 80 bp of flanking intronic sequences were synthesized (Thermo Fisher Scientific) and cloned in sense orientation using XhoI and NotI restriction sites of Exontrap vector pET01 (MoBiTec) to generate mini-genes that were validated by Sanger sequencing. The HEK293T cells were seeded in a 96-well plate at a cell density of 1.5 x 10^4^ and transfected the next day with 100 ng of mini-genes using Lipofectamine 3000 transfection reagent (Invitrogen), in 4 biological replicates. Cells were harvested 48 hrs post-transfection, and total RNA was extracted with QIACube using RNeasy kit with on-column DNase I treatment (Qiagen). For each sample, 0.5–1 μg of total RNA was converted into cDNA with SuperScript III reverse transcriptase (Invitrogen) and a vector-specific primer: 5’-AGGGGTGGACAGGGTAGTG-3’. Samples were diluted with water, and cDNA corresponding to 1.5 ng of RNA input was used for each qRT–PCR reaction. Splicing products of each mini-gene were amplified using a common primer pair F: 5’-CACCTTTGTGGTTCTCACTTGG-3’ and R: 5’-AGCACTGATCCACGATGCC-3’, corresponding to vector exons V1 and V2 (**Figure 4, Figure 5, Figure S6**). An assay with primers F: 5’-CCGTGACCTTCAGACCTTGG-3’ and R: 5’-AGAGAGCAGATGCTGGTGCA-3’targeting Exontrap vector exon V2 was used as a control. All PCR-amplified splicing products were resolved by agarose gel electrophoresis. Specific bands were cut out from the gel, purified and validated by Sanger sequencing. Co-transfection of E5 mini-gene with 10 SFs was performed in HEK293T cells for 48 hrs, followed by RNA extraction and analysis of splicing patterns. Similar analyses were performed for four select SFs (SRSF2, SRSF3, CELF1, and ELAVL2) in SW-780, HT-1376, and HTB-9 cells. The E5 construct was also transfected into a panel of 10 cell lines for 24 hrs, and splicing analysis was performed as described above.

### Modulation of *A3B* exon 5 splicing

Cell lines (HT-1376, HTB-9 and HeLa) were grown in 12-well plates at a density of 1.5 x 10^5^ cells/well and treated with DMSO or 1, 10, 20, 40, and 100 nM pladienolide B (Santa Cruz, sc-391691) reconstituted in DMSO, in biological triplicates per condition. Cells were harvested after 8, 16, 24, and 36 hrs of treatment separately for RNA and protein analysis. cDNA was synthesized from 300 ng of RNA in a 20 µl reaction using the RT^2^ First-Strand cDNA kit and random hexamers (Qiagen). cDNA corresponding to 30 ng of total RNA per reaction was used for PCR performed with sets of primers to amplify different fragments of *A3B*: primer pair 1 – F_ex1: 5’-GGACAGGGACAAGCGTATCT-3’, R_ex6: 5’-GCTCCAGGAGATGAACCAAG-3’; primer pair 2 – F_ex5: 5’-CCAGCACATGGGCTTTCTAT-3’, R_ex8: 5’-GAGATGGTGGTGAACGGTCT-3’ and primer pair 3 – F_ex4: 5’-GCCTTGGTACAAATTCGATGA-3’, R_ex6: 5’-TGTGTTCTCCTGAAGGAACG-3’. PCR-amplified splicing products were resolved by agarose gel electrophoresis, and specific bands were cut out from the gel, purified, and validated by Sanger sequencing. Quantification of *A3B1* and *A3B3* splicing products was performed with qRT-PCR using isoform-specific TaqMan assays (**Note S4**) and cDNA corresponding to 10 ng of total RNA per reaction, as previously described^1^. Expression of *A3B1* and *A3B3* was normalized by the expression of endogenous controls *GAPDH* (assay 4326317E) and *PPIA* (assay 4326316E). Water and genomic DNA were used as negative controls for all assays. Expression was measured as *C*_t_ values (PCR cycle at detection threshold) and calculated as ΔΔ*C*t in relation to control (DMSO) groups of samples. For protein analysis, whole-cell extracts were harvested in RIPA buffer with proteinase inhibitor (Fisher Scientific) and used for Western blotting as described in **Note S2**. Shown are representative results of one of three independent experiments.

### Cell-based cytosine deamination assay

To determine base substitution rates, we used an ssDNA oligo (**Figure S9A**), modified based on a previously reported oligo^66^. HT-1376 cells were nucleofected with 100 pmol of oligo alone or together with *A3B1* plasmid (program CM-130) with SF Cell Line 4D X Kit L (V4XC-2024) on Lonza 4D-Nucleofector. Nucleofected cells were then plated in a six-well plate and after 8 hrs, cells transfected only with oligo were treated with DMSO or pladienolide B for 64 hours. Cells were then lysed with QuickExtract DNA Extraction Solution (Lucigen) and PCR-amplified using primers: Forward, 5′-TGATGATGTGAGTGGTGGATGA-3′; Reverse, 5′-TCATCAACACCTACCACACAC-3′. PCR products were gel-purified with a DNA gel extraction kit (QIAquick Gel Extraction Kit, Qiagen) and used for library preparation with TruSeq/ChIP-Seq reagents (Illumina), to generate 75 bp paired-end sequencing reads. Samples were barcoded, multiplexed and subjected to deep sequencing on the Illumina MiSeq instrument. The FASTQ files were aligned with custom reference (sequence of ssDNA oligo) using the BWA-MEM algorithm and then indexed by SAMtools.

### siRNA knockdown of SF3B1

The HT-1376 cells were transfected with scrambled siRNA (#1022076), SF3B1 siRNA-1 (SI00715932), or siRNA-2 (SI04154647), all from Qiagen, using Lipofectamine RNAiMAX Reagent (ThermoFisher). Cells were harvested after 36 hrs for RNA and protein and analyzed for *A3B* exon 5 skipping by PCR as described above. SF3B1 knockdown was confirmed by Western blot with an anti-SF3B1 antibody (Abcam, ab172634, 1:1000 dilution), and GAPDH control as described above.

### Computational analysis

All data processing and analyses were performed using R package versions (3.2.4-3.4.0), SPSS version 25, and NIH High-Performance Computing Biowulf cluster.

## Data and reagents availability

The authors declare that data supporting the findings of this study are available from TCGA or within the paper and its supplementary information files. Additional information, protocols, and reagents can be provided on request to the corresponding author (LPO).

## Data deposition

The RNA-seq dataset for T47D cells infected with SeV and uninfected cells has been deposited to NCBI Short Read Archive (SRA), accession number PRJNA512015.

## URLs

Firehose Broad GDAC: https://gdac.broadinstitute.org/; Firebrowse: http://firebrowse.org/#; The Cancer Genome Atlas (TCGA): http://cancergenome.nih.gov; cBioPortal: http://www.cbioportal.org/index.do; Broad Institute Cancer Cell Line Encyclopedia (CCLE) https://portals.broadinstitute.org/ccle; Protein Data Bank (PDB): http://www.rcsb.org/pdb/home/home.do;

Clustal Omega: (http://www.ebi.ac.uk/Tools/msa/clustalo/); SpliceAid 2 (http://193.206.120.249/splicing_tissue.html); SFmap (http://sfmap.technion.ac.il/)

Human Splicing Finder (HSF), www.umd.be/HSF3/; The R project for statistical computing: http://www.r-project.org/; Integrative Genomics Viewer (IGV): http://www.broadinstitute.org/igv; ASpli R package: https://bioconductor.org/packages/release/bioc/vignettes/ASpli/inst/doc/ASpli.pdf

## Acknowledgments

The results presented here are in part based upon data generated by the TCGA Research Network. The study was supported by the Intramural Research Program of the National Cancer Institute - Division of Cancer Epidemiology and Genetics (L.P.O), AIDS Targeted Antiviral Program, Center for Cancer Research (V.K.P), and the NCI Innovation Award (R. B.). We thank the Cancer Genomics Research Laboratory of NCI for RNA-sequencing, Nathan Cole for help with retrieval of sliced RNA-seq BAM files from the NCI Data Commons portal and Dr. Marc-Henri Stern, Institute Curie, Paris, for critical comments and suggestions.

