## Supplementary materials for "Isoform-specific characterization implicates alternative splicing in *APOBEC3B* as a mechanism restricting APOBEC-mediated mutagenesis"

### Supplementary Materials, Banday et al


| Section details | Title | Page |
|---|---|---|
| **Tables** | | |
| Table S1 | Isoforms of the *APOBEC3* genes analyzed in TCGA samples | 2 |
| Table S2 | TCGA samples used in the analysis | 2-3 |
| Table S3 | Analysis of splicing factor (SF) binding sites within exons of *A3A* and *A3B* isoforms affected by alternative splicing | 4-5 |
| Table S4 | Expression constructs for alternative splicing factors used for co-expression studies | 6 |
| Table S5 | Summary for HSF analysis of branch point sites (BPS) within *A3B* introns | 6 |
| **Figures** | | |
| Figure S1 | Splicing patterns of *A3A* and *A3B* genes in TCGA tumor and adjacent normal tissues (N=11,058) based on RNA-seq reads for specific exon-exon junctions | 7 |
| Figure S2 | Evaluation of possible dominant-negative effects of catalytically inactive A3A and A3B isoforms | 8 |
| Figure S3 | Analysis of the activity of A3B protein isoforms using HIV-1 infectivity inhibition assays | 9 |
| Figure S4 | Distribution of APOBEC-signature mutations in TCGA samples | 10 |
| Figure S5 | Percent spliced-in index (PSI) for *A3B* alternative splicing isoforms in matched paired tumor and adjacent normal tissues in TCGA | 11 |
| Figure S6 | Analysis of alternative splicing in *A3A* and *A3B* with Exontrap assays | 12 |
| Figure S7 | Overexpression of SRSF2 increases the *A3B* exon 5 skipping in cancer cell lines | 13 |
| Figure S8 | Effects of pladienolide B treatment and siRNA-*SF3B1* knockdown on *A3B* exon 5 skipping in cancer cell lines | 14 |
| Figure S9 | Cell-based cytosine deamination assay | 15 |
| Figure S10 | Analysis of APOBEC-mediated mutagenesis in six TCGA cancers in relation to mutations in select splicing factors | 16 |
| **Notes** | | |
| S1 | Analysis of *A3A* and *A3B* transcripts based on exon-exon junctions and total expression using qRT-PCR and RNA-seq | 17-21 |
| S2 | Evaluation of the available antibodies for the detection of A3A and A3B protein isoforms | 22 |
| S3 | Purification/enrichment of recombinant A3A and A3B protein isoforms | 23 |
| S4 | Isoform-specific TaqMan expression assays for *A3B* transcripts | 24 |
| **Supplementary References** | | 25 |

**TABLES**

**Table S1. Isoforms of the *APOBEC3* genes analyzed in TCGA samples**

| Gene | Isoform ID in TCGA RNA-seq | UCSC ID in hg19 | Designation in the paper | Isoform type | Splicing event |
|---|---|---|---|---|---|
| A3A | uc003awn.2 | uc003awn.2 | A3A1 | Canonical | N/A |
| A3A | uc011aob.1 | uc011aob.1 | A3A2 | Alternative | Cryptic splice site in exon 2 |
| A3AB | uc011aoc.1 | uc011aoc.1 | A3AB | Deletion | N/A |
| A3B | uc003awo.1 | uc003awo.2 | A3B1 | Canonical | N/A |
| A3B | uc003awp.1 | uc003awp.2 | A3B2 | Alternative | Cryptic splice site in exon 6 |
| A3B | uc003awq.1 | uc003awq.2 | A3B3 | Alternative | Skipping of exon 5 |

**Table S2. TCGA samples used in the analysis**

| Number | Abbreviation | Cancer type | Samples, N[#] |
|---|---|---|---|
| 1 | ACC | Adrenocortical Carcinoma | 92 |
| 2 | BLCA | Bladder Urothelial Carcinoma | 412 |
| 3 | BRCA | Breast Invasive Carcinoma | 1098 |
| 4 | CESC | Cervical Squamous Cell Carcinoma and Endocervical Adenocarcinoma | 307 |
| 5 | CHOL | Cholangiocarcinoma | 51 |
| 6 | COAD | Colon Adenocarcinoma | 460 |
| 7 | DLBC | Lymphoid Neoplasm Diffuse Large B-cell Lymphoma | 58 |
| 8 | ESCA | Esophageal Carcinoma | 185 |
| 9 | GBM | Glioblastoma Multiforme | 613 |
| 10 | HNSC | Head and Neck Squamous Cell Carcinoma | 528 |
| 11 | KICH | Kidney Chromophobe | 113 |
| 12 | KIRC | Kidney Renal Clear Cell Carcinoma | 537 |
| 13 | KIRP | Kidney Renal Papillary Cell Carcinoma | 323 |
| 14 | LAML | Acute Myeloid Leukemia | 200 |
| 15 | LGG | Brain Lower Grade Glioma | 516 |
| 16 | LIHC | Liver Hepatocellular Carcinoma | 377 |
| 17 | LUAD | Lung Adenocarcinoma | 585 |
| 18 | LUSC | Lung Squamous Cell Carcinoma | 504 |
| 19 | MESO | Mesothelioma | 87 |
| 20 | OV | Ovarian Serous Cystadenocarcinoma | 602 |
| 21 | PAAD | Pancreatic Adenocarcinoma | 185 |
| 22 | PCPG | Pheochromocytoma and Paraganglioma | 179 |
| 23 | PRAD | Prostate Adenocarcinoma | 499 |
| 24 | READ | Rectum Adenocarcinoma | 171 |
| 25 | SARC | Sarcoma | 261 |
| 26 | SKCM | Skin Cutaneous Melanoma | 470 |
| 27 | STAD | Stomach Adenocarcinoma | 443 |
| 28 | TGCT | Testicular Germ Cell Tumors | 150 |
| 29 | THCA | Thyroid Adenocarcinoma | 503 |
| 30 | THYM | Thymoma | 124 |
| 31 | UCEC | Uterine Corpus Endometrioid Carcinoma | 560 |
| 32 | UCS | Uterine Carcinosarcoma | 57 |
| 33 | OVM | Uveal Melanoma | 80 |
| Total | | | 11330 |

### Numbers of samples used for specific analyses may vary based on the availability of data for RNA-seq, APOBEC-signature mutations, and other variables.

**Table S3. Analysis of splicing factor (SF) binding sites within exons of *A3B* isoforms affected by alternative splicing**

| Isoform | Type of alternative splicing | Splicing event | The exonic sequence for SFs prediction, bp | SF | Another name | Target site, bp position within query sequence |
|---|---|---|---|---|---|---|
| **A3B2** | Between exon 5 and cryptic acceptor splice site in exon 6 | Loss of 75 bp in exon 6 | 1-ggctaagaatcttctctgtggcttttacggccgccatgcggagctgcgcttcttggacctggttccttctttgca-75 | CUG-BP | CELF1 | UGCGG (37-41), AGCUG (42-46) |
| | | | | HuB | ELAVL2 | CUUUUA (22-27) |
| | | | | hnRNP A1 | HNRNPA1 | UACGGC (26-31) |
| | | | | hnRNPF | HNRNPF | GUGGCUU (18-24) |
| | | | | hnRNP H1 | HNRNPH1 | AAGAA (5-9) |
| | | | | hnRNP H2 | HNRNPH2 | AUCUUC (5-9) |
| | | | | hnRNPM | HNRNPM | GGUUCCUU (61-68) |
| | | | | Sam68 | KHDRBS1 | UUUUAC (23-28) |
| | | | | MBNL1 | MBNL1 | GGCUUU (20-25, 46-51) |
| | | | | hnRNP I (PTB) | PTBP1 | AUCUUC (10-14), UCUUCUCU (10-17), UUCUUG (50-55), UUCUUU (67-72), UCUUU (68-72) |
| | | | | SC35 | SRSF2 | GGCCGCCA (29-36), GGUUCCUU (61-68) |
| | | | | SRp40 | SRSF5 | ACGGC (27-31), GCUGC (43-47) |
| | | | | SRp55 | SRSF6 | UGCGGA (37-42) |
| | | | | SRp30c | SRSF9 | CGGAG (39-43), UGGAC (54-58) |
| | | | | TIA-1 | TIA1 | CUUUUA (22-27), CUUUG (69-73) |
| | | | | TIAL1 | TIAL1 | CUUUUA (22-27), CUUUG (69-73) |
| | | | | HTra2alpha | TRA2A | AAGAA (5-9) |
| | | | | HTra2beta1 | TRA2B | AAGAA (5-9) |
| **A3B3** | Between exon 4 and exon 6 | Loss of entire exon 5, 129 bp | 1-gatacctgatggatccagacacattcactttcaacttgatacctgatggatccagacacattcactttcaacttcctggacaatggcacctgggtcctgatggaccagcacatgggcttt | CUG-BP | CELF1 | UGAUG (44-48), UCCUG (74-78), UCCUG (95-99) |
| | | | | HuB | ELAVL2 | CUUUC (28-32), CUUUC (65-69), CUUUC (117-121) |
| | | | | hnRNP F | HNRNPF | UGGGC (113-117) |
| | | | | hnRNP H1 | HNRNPH1 | UGGGU (91-95), UGGGC (113-117) |
| | | | | hnRNP H2 | HNRNPH2 | UGGGU (91-95), UGGGC (113-117) |

| | | | | | | |
|---|---|---|---|---|---|---|
| | | | ctatgcaac-129 | hnRNP H3 | HNRNPH3 | UGGGU (91-95), UGGGC (113-117) |
| | | | | KSRP | KHSRP | UGGGU (91-95) |
| | | | | MBNL1 | MBNL1 | GGCUUU (115-120) |
| | | | | Nova-1 | NOVA1 | UUCAAC (30-35), UUCAAC (67-72) |
| | | | | Nova-2 | NOVA2 | UUCAAC (30-35), UUCAAC (67-72) |
| | | | | SC35 | SRSF2 | UCAAC (31-35), UCAAC (68-72), ACUUC (71) |
| | | | | SRp20 | SRSF3 | GAUC (12-15), UUCAC (24-28), GAUC (49-52), UUCAC (61-65) |
| | | | | SRp40 | SRSF5 | UUCCUGG (73-79) |
| | | | | 9G8 | SRSF7 | UGGACAA (77-83) |
| | | | | SRp30c | SRSF9 | UGGAU (10-14), UGGAU (47-51), UGGAC (77-81), UGGAC (101-105) |
| | | | | TIA-1 | TIA1 | CUUUC (28-32), CUUUC (65-69), CUUUC (117-121) |
| | | | | TIAL1 | TIAL1 | CUUUC (28-32), CUUUC (65-69), CUUUC (117-121) |
| | | | | HTra2beta1 | TRA2B | UCAAC (31-35), UCAAC (68-72) |
| | | | | YB-1 | YBX1 | GAUC (12-15), GAUC (49-52), CACC (87-90) |

Bioinformatic predictions were performed with SFmap [1] and SpliceAid 2 [2]

**Table S4. Expression constructs for alternative splicing factors used for co-expression studies**

| Construct | OriGene* catalog ID | Construct | NCBI ID | Matching UCSC Isoforms, hg19 |
|---|---|---|---|---|
| 1 | RC206275 | CELF1-T1 | NM_006560 | uc001nfn.3 |
| 2 | RC222434 | CELF1-T3 | NM_001025596 | uc001nfp.3, uc001nfl.3, uc001nfr.1 |
| 3 | RC230208 | CELF1-T4 | NM_001172639 | uc001nfk.2 |
| 4 | RC229918 | ELAV2 | NM_001171195 | uc003zpt.3 |
| 5 | RC223971 | TIAL1 | NM_003252 | uc001lei.1 |
| 6 | RC200263 | KHDRBS1 | NM_006559 | uc001buc.2, uc001bub.4 |
| 7 | RC209660 | SFRS3 (SRSF3) | NM_003017 | uc003omj.3, uc011dtp.1 |
| 8 | RC209842 | SFRS2 (SRSF2) | NM_003016 | uc002jsv.3, uc010wtg.2 |
| 9 | RC203314 | HNRNPA1 | NM_002136 | uc001sfo.3, uc009znj.1, uc001sfm.3, uc001sfn.3 |
| 10 | RC219704 | MBNL1 | NM_207295 | uc003ezn.3 |

*expression constructs were purchased from OriGene

**Table S5. Summary for HSF analysis of branch point sites (BPS) within *A3B* introns**

| Intron of *A3B* | Position of the strongest motif, bp upstream of exon | BPS motif | Consensus score (0-100) for a top BPS in each intron | Number of potential BPS with a score above threshold (>67) as per HSF |
|---|---|---|---|---|
| 1 | -58 | ccccgAg | 83.31 | 3 |
| 2 | -37 | ctctcAg | 93.91 | 6 |
| 3 | -30 | gcctgAc | 93.44 | 3 |
| 4 | -38 | gtcttAg | 80.45 | 2 |
| 5 | -26 | acctcAc | 95.75 | 10 |
| 6 | -77 | tcctgAg | 90.81 | 6 |
| 7 | -40 | ctctcAc | 96.63 | 7 |

HSF, Human Splicing Finder – www.umd.be/HSF3/

**FIGURES**

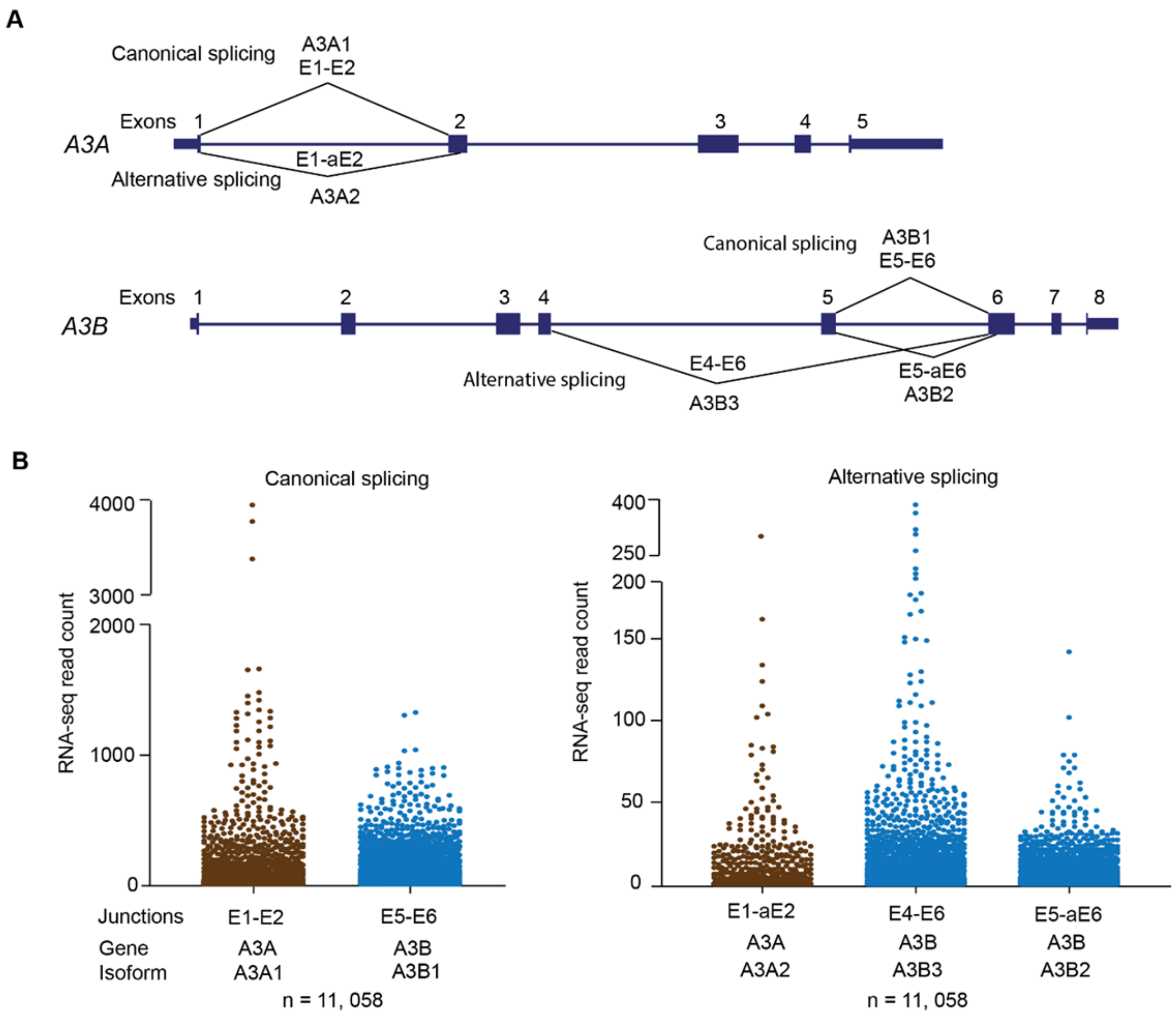


**Figure S1. Splicing patterns of *A3A* and *A3B* genes in TCGA tumor and adjacent normal tissues (N=11,058) based on RNA-seq reads for specific exon-exon junctions. A)** Two alternatively spliced isoforms were observed for *A3A* - the canonical isoform *A3A1* defined by E1-E2 junction, and the alternative *A3A2* isoform defined by E1-aE2 junction through a cryptic acceptor site in exon 2 (aE2). Three alternatively spliced isoforms were observed for *A3B* - the canonical isoform *A3B1* defined by E5-E6 junction, and the alternative isoforms *A3B2* and *A3B3* defined by the usage of an alternative acceptor site (E5-aE6 junction), and exon 5 skipping (E4-E6 junction), respectively. **B**) Counts of RNA-seq reads supporting the canonical exon junctions E1-E2 and E5-E6 for *A3A1* and *A3B1* isoforms, respectively. **C**). Counts of RNA-seq reads supporting the alternative exon junctions E1-aE2 for *A3A*2, E5-aE6 for *A3B2* and E4-E6 for *A3B3*.

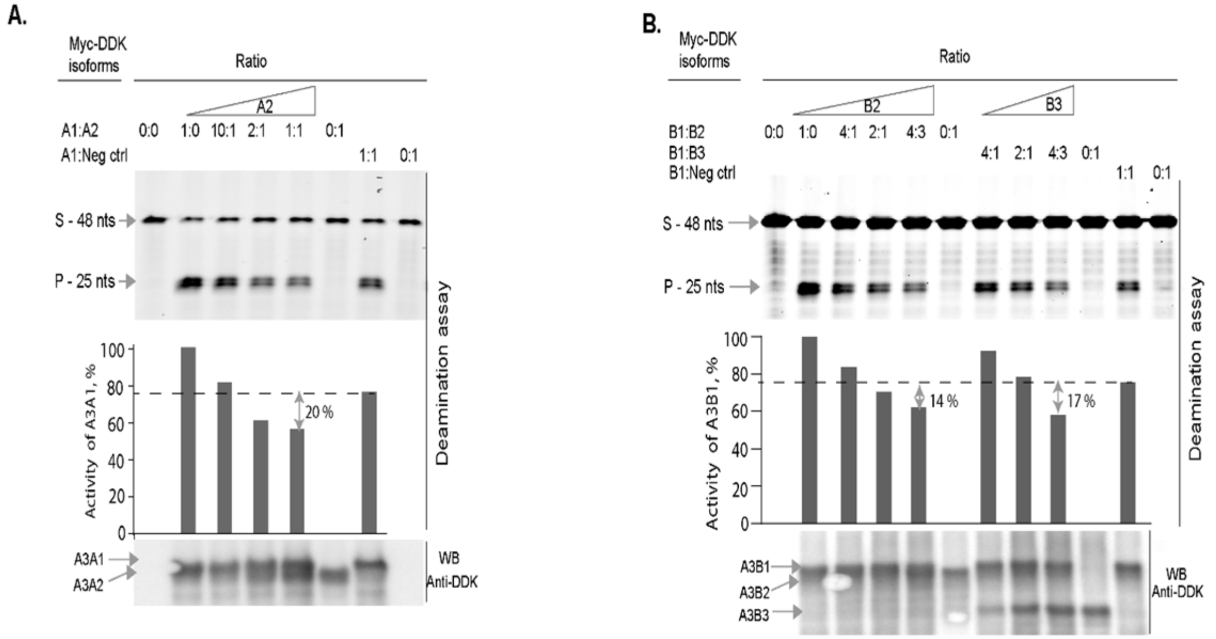


**Figure S2. Evaluation of possible dominant-negative effects of catalytically inactive A3A and A3B isoforms. A)** Deaminating activity of the mutagenic protein isoform A3A1 is moderately inhibited (by 20%) by the corresponding non-mutagenic A3A2 protein isoform. **B**) Deaminating activity of the mutagenic isoform A3B1 is moderately inhibited (by 14 and 17%) by the corresponding non-mutagenic A3B2 and A3B3 protein isoforms, respectively. All assays were performed using enriched fractions of semi-purified proteins (**Note S3**). The amounts of mutagenic isoforms and total reaction volumes were kept constant while the amounts of non-mutagenic isoforms were increased. The deamination activity was compared to the mutagenic isoforms alone, and to the mutagenic isoforms mixed in a 1:1 ratio with a protein extracted from lysates of untransfected cells to account for inhibition caused by non-specific endogenous proteins. Results are representative of four independent experiments.

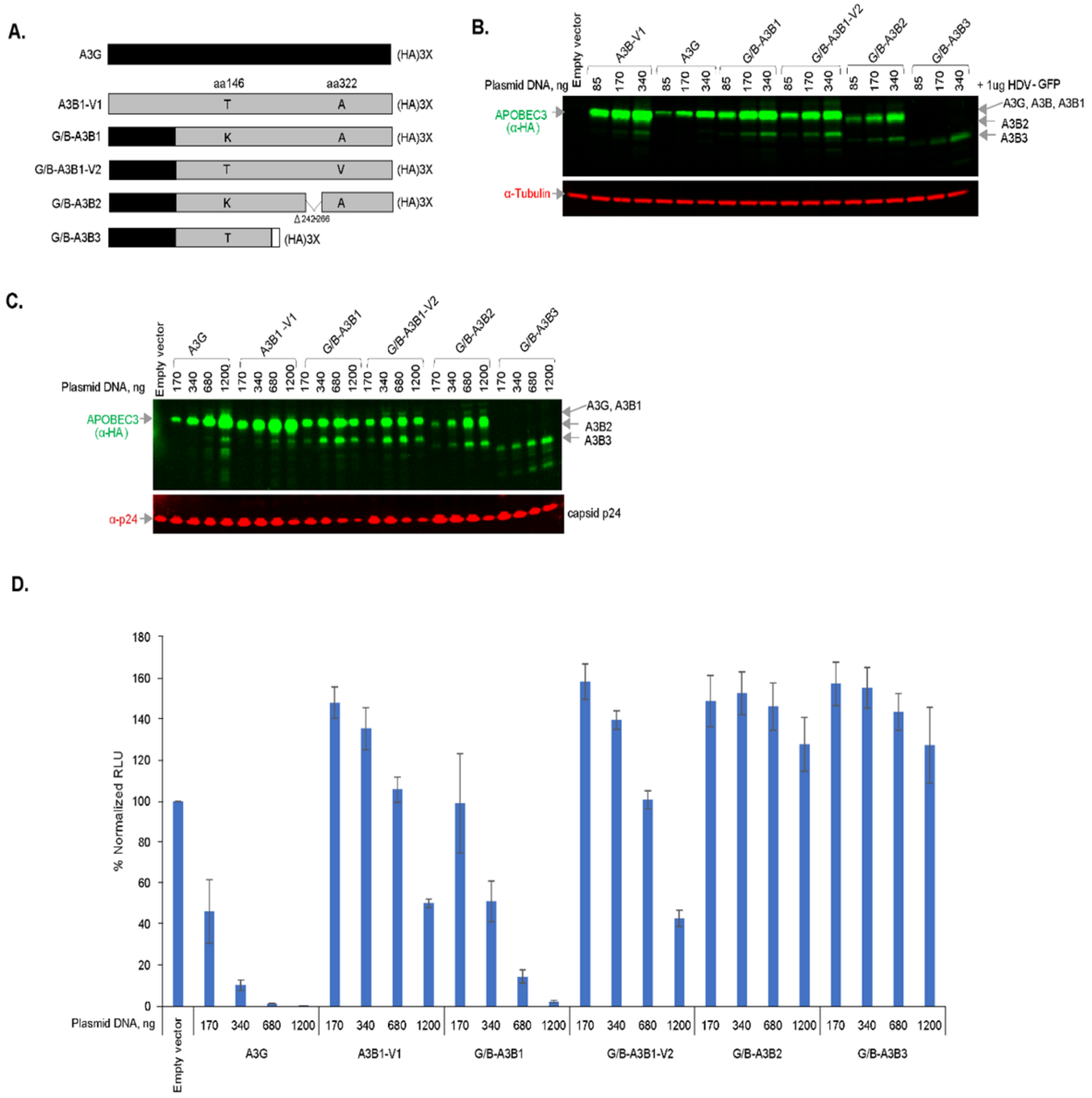


**Figure S3. Analysis of the activity of A3B protein isoforms using HIV-1 infectivity inhibition assays. A)** Fusion constructs were generated by replacing 63 aa of the N-terminal region of A3B with a similar region of APOBEC3G (A3G). **B**) All G/B fusion constructs were transfected into HEK293T cells; protein expression and stability were established as G/B-A3B1 = G/B-A3B1-V2 > G/B-A3B2 >>> G/B-A3B3. **C**) Western blotting is showing the viral packaging of recombinant proteins. Consistent with their steady-state expression levels, G/B-A3B2 and G/B-A3B3 fusions were incorporated into virions to a lesser extent than G/B-A3B1 and G/B-A3B2-V2. **D**) A3B2 and A3B3 proteins had no detectable effect on the inhibition of HIV-1 infectivity. As expected based on previous reports [3,4], a fusion with the A3G N-terminal region enhanced the antiviral activity of A3B1 (compare A3B1-V1 with G/B-A3B1). A3B1-V1 inhibited HIV-1 infectivity in a dose-dependent manner, but A3B2 and A3B3 isoforms did not. Shown are means ± SD of biological triplicates.

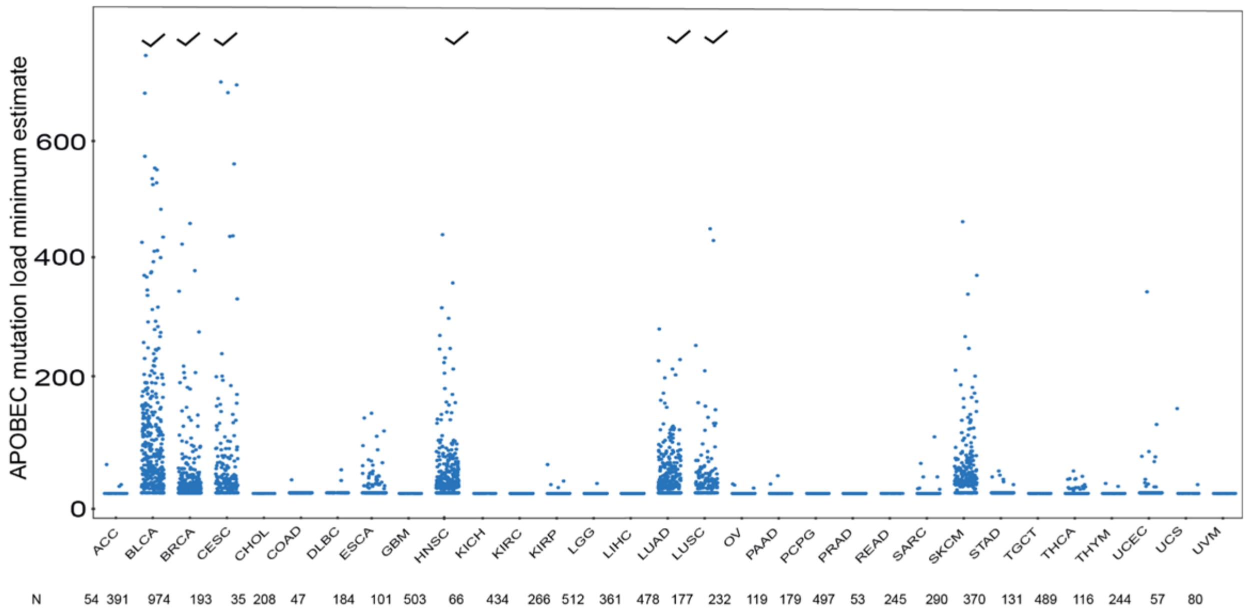


**Figure S4. Distribution of APOBEC-signature mutations in TCGA samples.** Abbreviations for cancer types are explained in **Table S2**; each dot represents a sample. N - the number of samples with mutations for each cancer type. Tick marks indicate cancer types with the highest numbers of samples with APOBEC-signature mutations; a set of 6 cancer types (BLCA, BRCA, CESC, HNSC, LUAD, and LUSC) was selected for further analysis in relation to the expression of the mutagenic isoforms *A3A1* and *A3B1* (**Figure 3**). Although SKCM is enriched in samples with APOBEC-signature mutations, this cancer type was not analyzed because of the overlap between mutations caused by APOBECs and UV radiation[5].

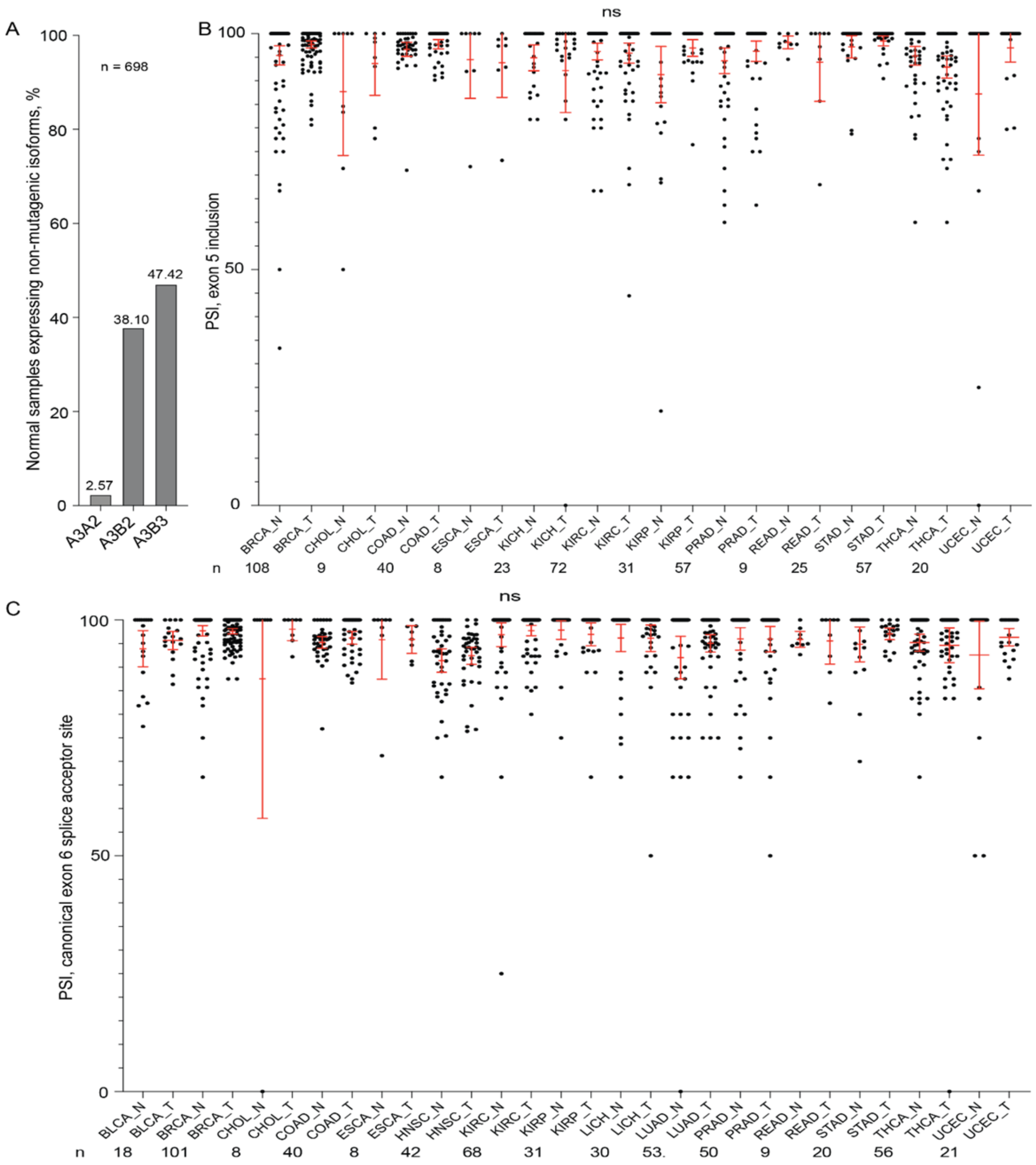


**Figure S5. Percent spliced-in index (PSI) for *A3B* alternative splicing isoforms in matched paired tumor and adjacent normal tissues in TCGA. A)** Percentage of matched normal samples expressing alternative isoforms of *A3A* and *A3B* with ≥1 RNA-seq reads for unique exon-exon junctions. **B** and **C**) PSI values for the canonical splicing isoforms - exon 5 inclusion and splicing through the canonical exon 6 acceptor site (E5-E6 junction) in several cancer types. These cancers show no significant differences (indicated as "ns") in PSI levels between matched paired tumor (T) and normal (N) samples. The cancer types with significant differences in PSI levels between paired tumor and normal samples are shown in **Figure 4**.

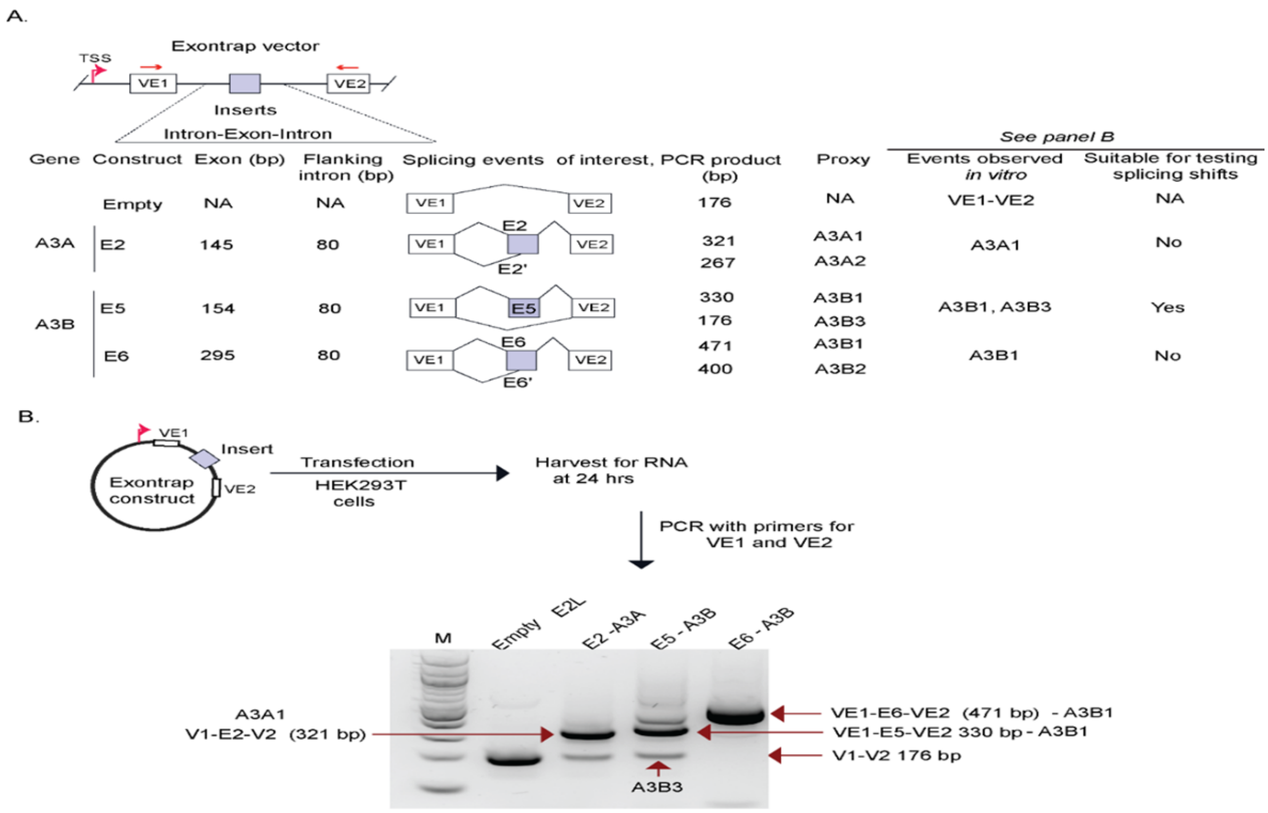


**Figure S6. Analysis of alternative splicing in *A3A* and *A3B* with Exontrap assays. A)** Schematic representation of the Exontrap strategy - the alternatively spliced exons (E) of *A3A* (exon 2) and *A3B* (exons 5 and 6) were cloned into an Exontrap vector, which carries a transcription start site (TSS) and vector exons VE1 and VE2. DNA fragments are cloned into insert position and tested for exon inclusion (trapping) between vector exons VE1 and VE2. Due to the large size of flanking introns (>1Kb), constructs were generated for each alternative exon with only 80 bp flanking intronic regions, assuming these sequences include essential cis-elements such as splice and branch point sites. The goal of the initial screening was to identify Exontrap constructs that generate predicted splicing events representing all *A3A* and *A3B* isoforms. A proxy for *A3A1* is represented by splicing of vector exon VE1 with the insert exon E2 and then with vector exon VE2, as VE1- E2-VE2, while a proxy for *A3A1* is represented by splicing of VE1 with E2 through a cryptic internal splicing site (E2'), as VE1- E2'-VE2. Proxies for *A3B1* are represented by inclusion of E5 or E6, as VE1- E5-VE2 or VE1- E6-VE2; a proxy for *A3B2* is represented by splicing of VE1 with E6 through a cryptic splice site (E6'), as VE1- E6'-VE2, and a proxy for *A3B3* in this context is represented by exclusion of E5, as VE1-VE2.

**B**) Screening of Exontrap constructs transiently transfected into HEK293T cells for 24 hours. RNA was extracted, and PCR products were generated from cDNA with primers corresponding to vector exons VE1 and VE2 and resolved by agarose gel electrophoresis. Each band was cut out from the gel, purified and Sanger-sequenced to confirm the identity of splicing events. In HEK293T cells, only the E5 construct showed both splicing events of interest (representing *A3B1* and *A3B3* transcripts) and was selected for further detailed analysis. M – 100 bp size marker.

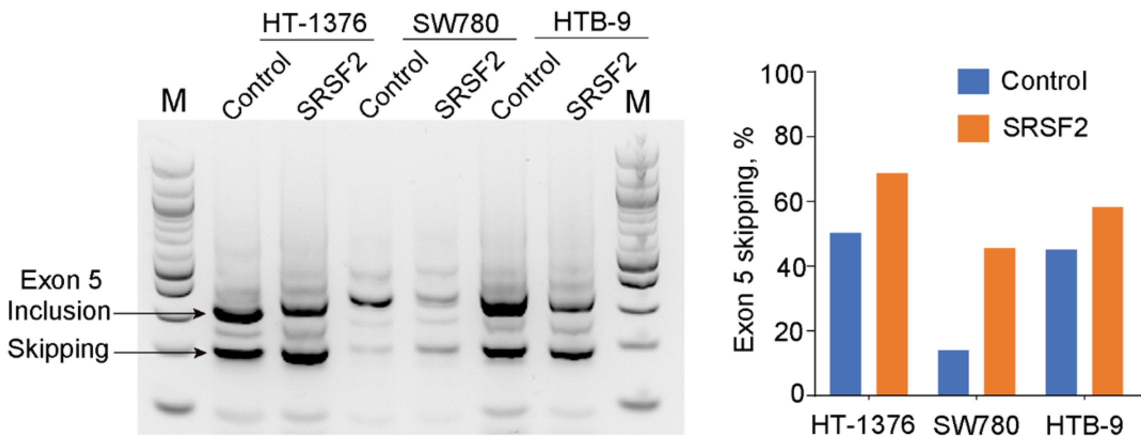


**Figure S7. Overexpression of SRSF2 increases the *A3B* exon 5 skipping in cancer cell lines.** Exontrap mini-gene E5 with exon 5 of *A3B* was transiently transfected into bladder cancer cell lines – HT-1376, SW780, and HTB-9 together with the *SRSF2* expression construct or alone (control). Cells were harvested after 24 hrs and splicing products corresponding to exon 5 inclusion (*A3B1*-type transcript) vs. skipping (*A3B3*-type transcript) were analyzed by RT-PCR and agarose gel electrophoresis. Band intensity was quantified using Image Lab software (BioRad) and plotted as % of exon 5 skipping of the combined expression of the *A3B1* and *A3B3* transcripts in each sample. Shown are results of one of the three independent experiments.

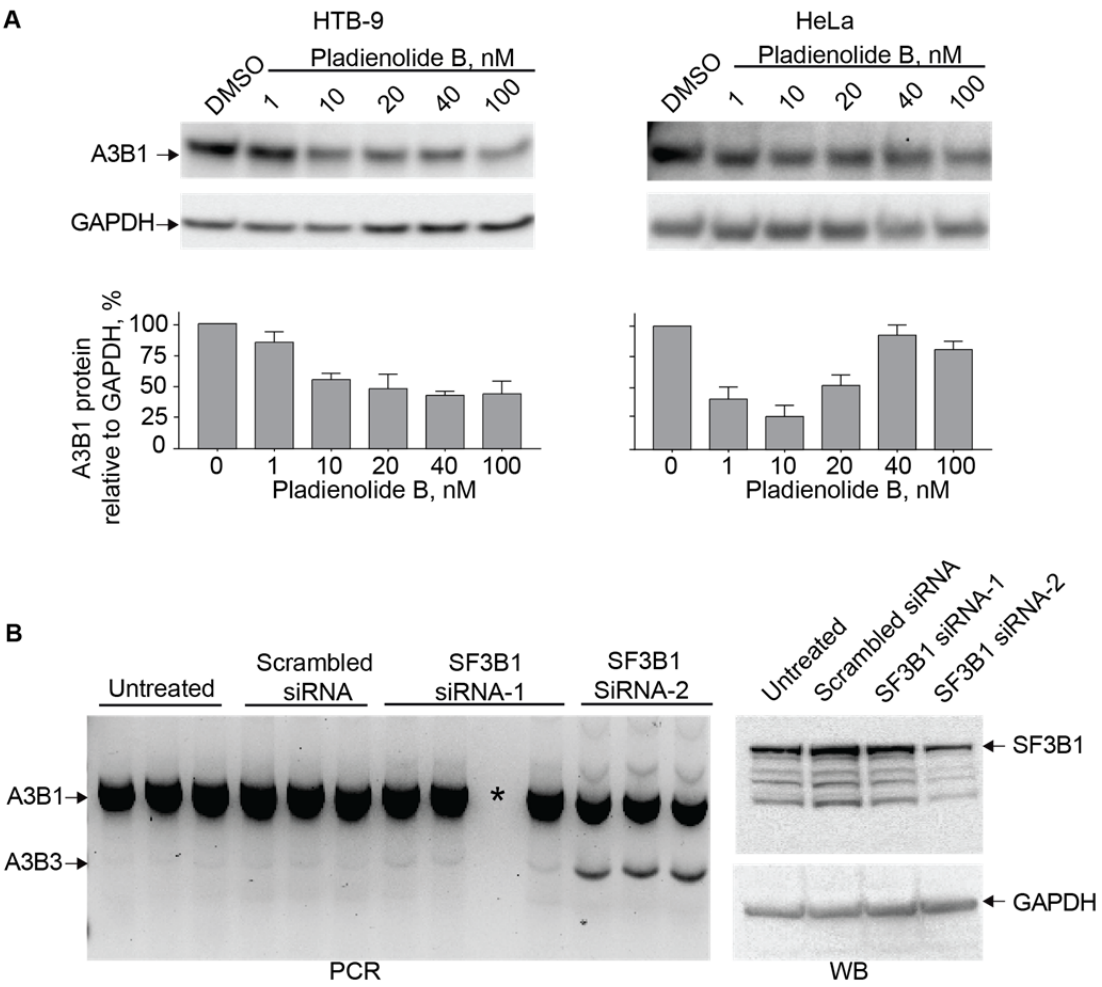


**Figure S8. Effects of pladienolide B treatment and siRNA-*SF3B1* knockdown on *A3B* exon 5 skipping in cancer cell lines. A)** A3B1 protein levels in HTB-9 and HeLa cells treated with pladienolide B for 36 hrs. Western blot (WB) analysis shows 20-70% reduction of A3B1 protein levels in cells treated with different concentrations of pladienolide B. Protein levels were determined by densitometry (ImageJ) and graphed as a percentage compared to A3B1 expression levels in the cells treated with DMSO (vehicle), after normalization by GAPDH expression (loading control). Shown are means ± SD of biological triplicates. The effect is observed only in HTB-9 but not in HeLa cells. **B)** siRNA-*SF3B1* knockdown in HT-1376 cells. Left panel: RT-PCR products analyzed by agarose gel electrophoresis showing the change in *A3B* exon 5 skipping in cells transfected for 36 hrs with *SF3B1* siRNA-1 or 2 comparing to untransfected cells and cells treated with scrambled siRNA (negative control), all in biological triplicates. Right panel: WB analysis with an anti-SF3B1 antibody confirms the S3FB1 protein knockdown, which was more successful using siRNA-2, judging from the RT-PCR results for *A3B1* exon 5 skipping and WB analysis for SF3B1 expression.* - empty lane.

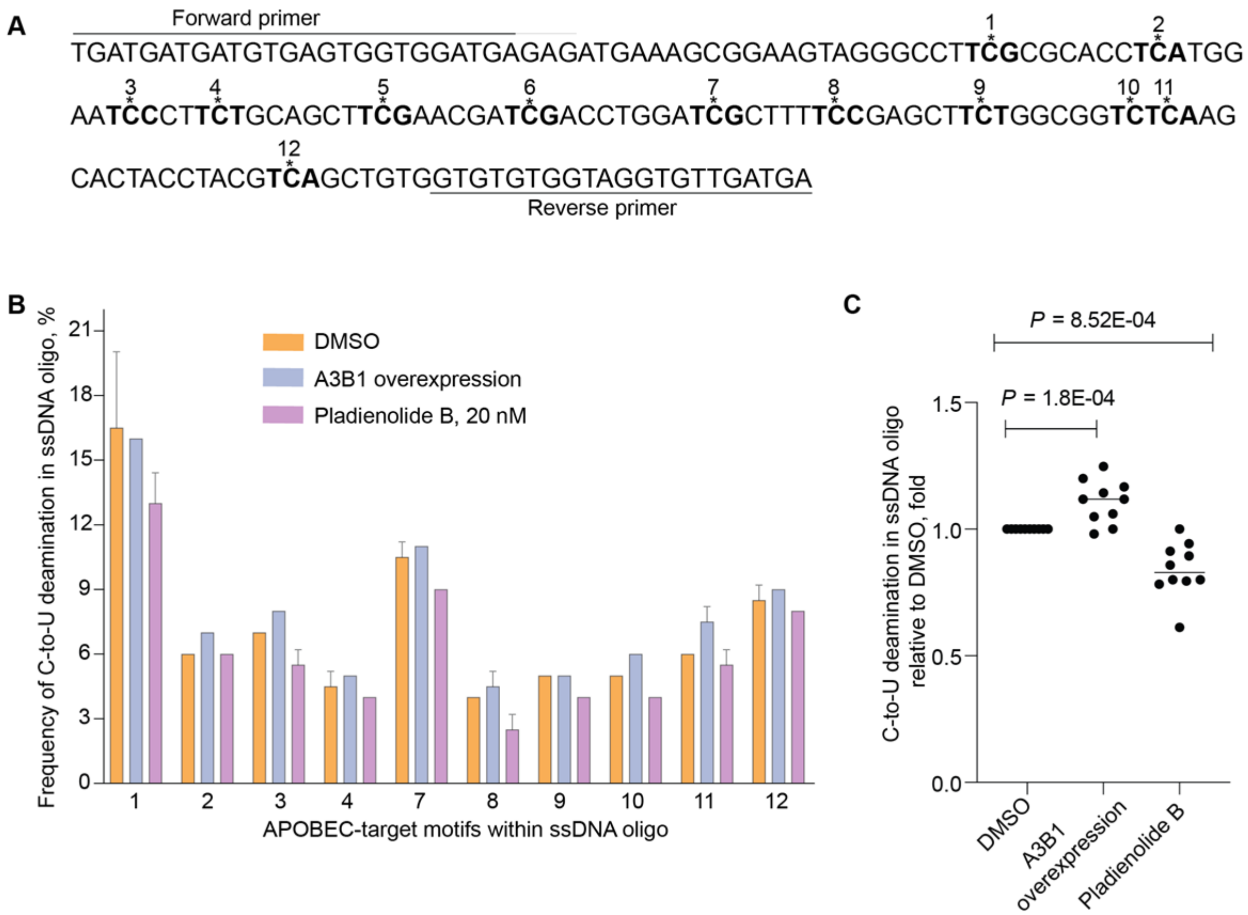


**Figure S9. Cell-based cytosine deamination assay.**

**A)** The sequence of the 169-bp ssDNA oligo. APOBEC sequence motifs are in bold with target cytosines indicated by asterisks and motif numbers (1-12). **B)** Frequencies (%) of cytosine deamination (C->U) within each APOBEC motif in different experimental conditions. Bladder cancer HT-1376 cells were nucleofected with the ssDNA oligo alone or together with *A3B1*-expressing plasmid (positive control); after 8 hours, cells transfected only with ssDNA oligo were treated with DMSO or pladienolide B for 64 hours. The ssDNA oligos were recovered from all the cells, PCR-amplified and deep-sequenced (see Materials and Methods). Shown are means ± SD of biological duplicates. APOBEC motifs #5 and #6 were excluded from the analysis because of low coverage of this region by the short (76 bp) paired sequencing reads not reaching the middle of the 169-bp PCR product. **C**) The ratios of cytosine deamination within all APOBEC motifs in the ssDNA oligo in indicated experimental conditions compared to DMSO-treated cells. Each dot represents the ratio of C->U deamination at one of the ten APOBEC-target motifs. Overall, these ratios were significantly higher in A3B1-overexpressing cells and lower in pladienolide B - treated cells compared to DMSO-treated control cells. *P*-values are for the Student’s t-test.

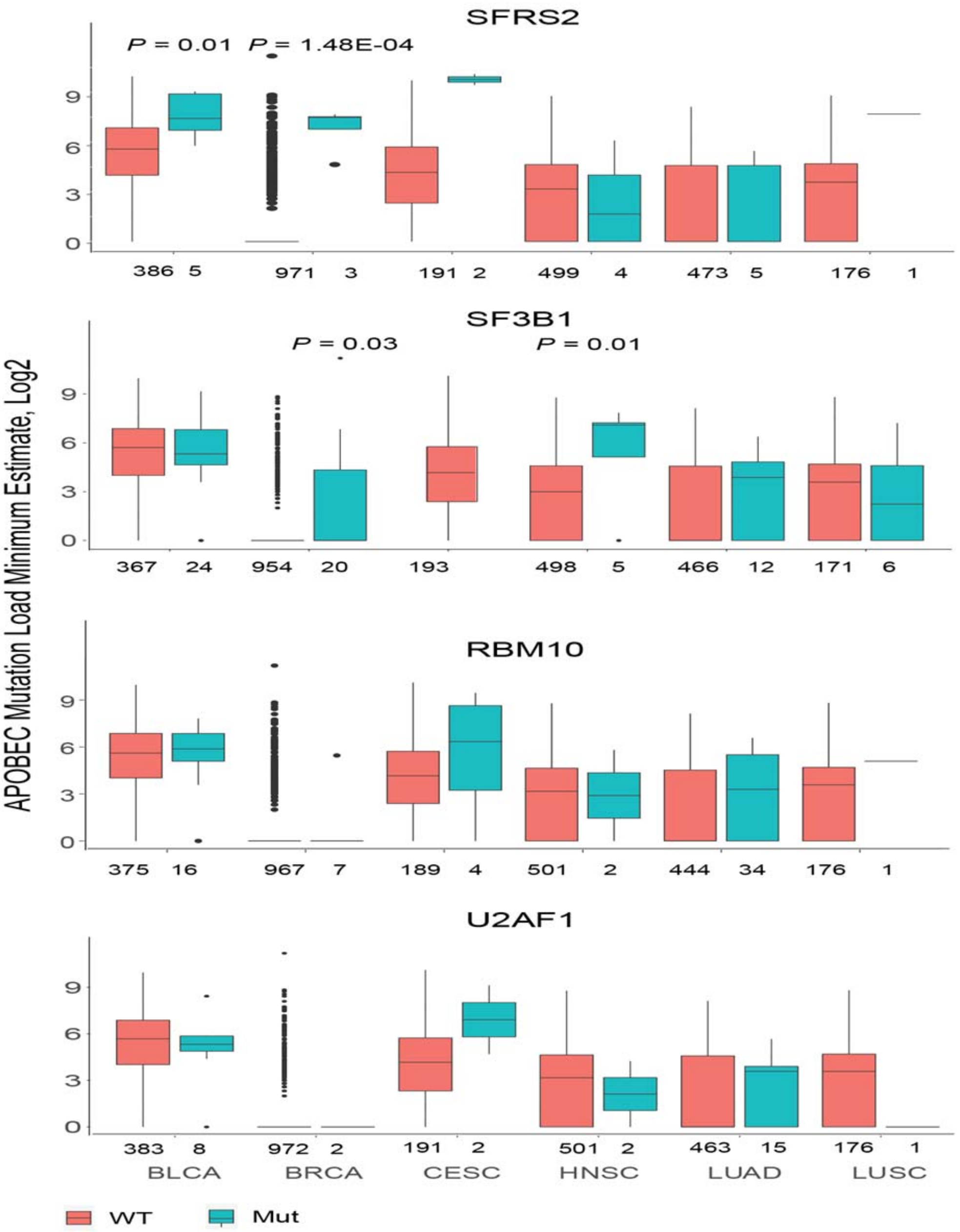


**Figure S10. Analysis of APOBEC-mediated mutagenesis in six TCGA cancers in relation to mutations in select splicing factors.** The analysis included nonsense, missense, splice site, and in-frame deletion/insertion mutations in splicing factors *SFRS2*, *SF3B1*, *RBM10,* and *U2AF1*, considering only driver mutations[6]. A trend for a higher load of APOBEC-signature mutations is observed in some cancer types with mutations in *SFRS2* and *SF3B1*.

#### NOTES

**Note S1. Analysis of *A3A* and *A3B* transcripts based on exon-exon junctions and total expression using qRT-PCR and RNA-seq**

```
A3A    ------------------------------------------------------------    0
A3B    ATGAATCCACAGATCAGAAATCCGATGGAGCGGATGTATCGAGACACATTCTACGACAAC    60


A3A    ------------------------------------------------------------    0
A3B    TTTGAAAACGAACCCATCCTCTATGGTCGGAGCTACACTTGGCTGTGCTATGAAGTGAAA    120


A3A    ------------------------------------------------------------    0
A3B    ATAAAGAGGGGCCGCTCAAATCTCCTTTGGGACACAGGGGTCTTTCGAGGCCAGGTGTAT    180


A3A    ------------------------------------------------------------    0
A3B    TTCAAGCCTCAGTACCACGCAGAAATGTGCTTCCTCTCTTGGTTCTGTGGCAACCAGCTG    240


A3A    ------------------------------------------------------------    0
A3B    CCTGCTTACAAGTGTTTCCAGATCACCTGGTTTGTATCCTGGACCCCCTGCCCGGACTGT    300


A3A    ------------------------------------------------------------    0
A3B    GTGGCGAAGCTGGCCGAATTCCTGTCTGAGCACCCCAATGTCACCCTGACCATCTCTGCC    360


A3A    ------------------------------------------------------------    0
A3B    GCCCGCCTCTACTACTACTGGGAAAGAGATTACCGAAGGGCGCTCTGCAGGCTGAGTCAG    420


A3A    ------------------------------------------------------------    0
A3B    GCAGGAGCCCGCGTGACGATCATGGACTATGAAGAATTTGCATACTGCTGGGAAAACTTT    480


A3A    ------------------------------------------------------------    0
A3B    GTGTACAATGAAGGTCAGCAATTCATGCCTTGGTACAAATTCGATGAAAATTATGCATTC    540

                                   78.8.3% identity between A3A-Ex2 and A3B-Ex5
A3A    ATGGAAGCCAGCCCAGCATCCGGGCCCAGACACTTGATGGATCCACACATATTCACTTCC    60
A3B    CTGCACCGCACGCTAAAGGAGATTCTCAGATACCTGATGGATCCAGACACATTCACTTTC    600
        ** *   **  * *         * **** ** *********** *** ******** *

A3A    AACTTTAACAATG---------GCATTGGAAGGCATAAGACCTACCTGTGCTACGAAGTG    111
A3B    AACTTTAATAATGACCCTTTGGTCCTTCGACGGCGCCAGACCTACTTGTGCTATGAGGTG    660
       ******** ****          * ** ** ***   ******** ******* ** ***

A3A    GAGCGCCTGGACAATGGCACCTCGGTCAAGATGGACCAGCACAGGGGCTTTCTACACAAC    171
A3B    GAGCGCCTGGACAATGGCACCTGGGTCCTGATGGACCAGCACATGGGCTTTCTATGCAAC    720
       ********************** ****  ************** **********  ****

        99.3% identity between A3A-Ex3 and A3B-Ex6
A3A    CAGGCTAAGAATCTTCTCTGTGGCTTTTACGGCCGCCATGCGGAGCTGCGCTTCTTGGAC    231
A3B    GAGGCTAAGAATCTTCTCTGTGGCTTTTACGGCCGCCATGCGGAGCTGCGCTTCTTGGAC    780
        ***********************************************************

A3A    CTGGTTCCTTCTTTGCAGTTGGACCCGGCCCAGATCTACAGGGTCACTTGGTTCATCTCC    291
A3B    CTGGTTCCTTCTTTGCAGTTGGACCCGGCCCAGATCTACAGGGTCACTTGGTTCATCTCC    840
```

```
         ************************************************************

A3A      TGGAGCCCCTGCTTCTCCTGGGGCTGTGCCGGGGAAGTGCGTGCGTTCCTTCAGGAGAAC        351
A3B      TGGAGCCCCTGCTTCTCCTGGGGCTGTGCCGGGGAAGTGCGTGCGTTCCTTCAGGAGAAC        900
         ************************************************************

A3A      ACACACGTGAGACTGCGTATCTTCGCTGCCCGCATCTATGATTACGACCCCCTATATAAG        411
A3B      ACACACGTGAGACTGCGCATCTTCGCTGCCCGCATCTATGATTACGACCCCCTATATAAG        960
         ***************** ******************************************

A3A      GAGGCACTGCAAATGCTGCGGGATGCTGGGGCCCAAGTCTCCATCATGACCTACGATGAA        471
A3B      GAGGCGCTGCAAATGCTGCGGGATGCTGGGGCCCAAGTCTCCATCATGACCTACGATGAG        1020
         ***** ***************************************************** 

         94.8% identity between A3A-Ex4 and A3B-Ex7
A3A      TTTAAGCACTGCTGGACACCTTTGTGGACCACCAGGGATGTCCCTTCCAGCCCTGGGAT        531
A3B      TTTGAGTACTGCTGGACACCTTTGTGTACCGCCAGGGATGTCCCTTCCAGCCCTGGGAT        1080
         *** ** ******************* *** *****************************

A3A      GGACTAGATGAGCACAGCCAAGCCCTGAGTGGGAGGCTGCGGGCCATTCTCCAGAATCAG        591
A3B      GGACTAGAGGAGCACAGCCAAGCCCTGAGTGGGAGGCTGCGGGCCATTCTCCAGAATCAG        1140
         ******** ***************************************************

A3A      GGAAACTGA       600
A3B      GGAAACTGA       1149
         *********
```

**I). High homology between coding sequences of *A3A* and *A3B* genes**

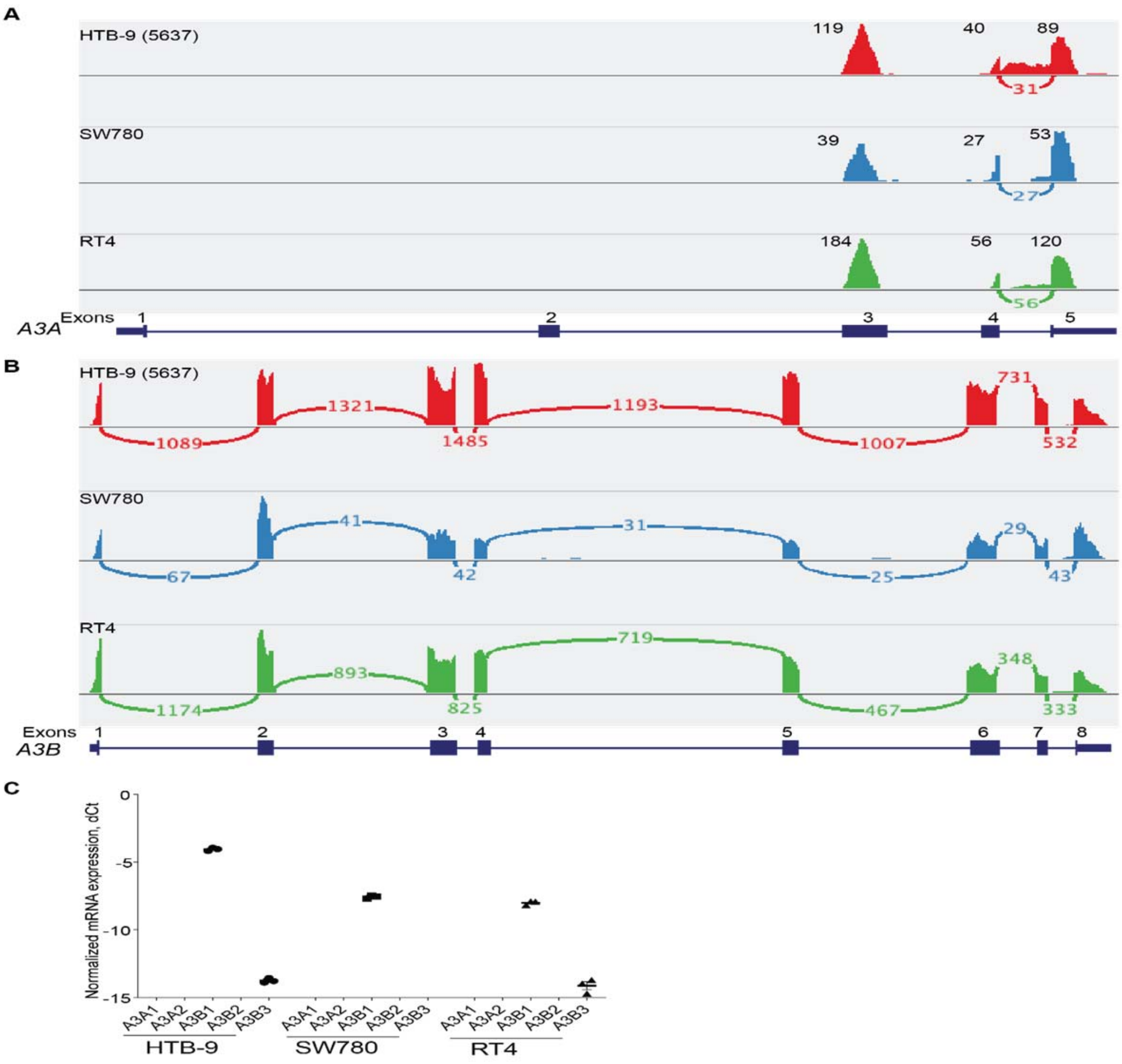


**II). Due to high sequence homology, *A3B* RNA-seq reads misalign to *A3A***

Expression of *A3A* and *A3B* genes visualized as IGV sashimi plots based on RNA-seq BAM files for bladder cancer cell lines HTB-9, SW780, and RT4 (from Broad Institute Cancer Cell Line Encyclopedia, CCLE). **A**) Within *A3A*, RNA-seq reads align only to exons 3-5; **B**) Within *A3B*, RNA-seq reads align to all exons. **C**) qRT-PCR analysis in corresponding cell lines shows high expression of the *A3B1* isoform and weak expression of the *A3B3* isoform, while no expression of the *A3A* isoforms. When expression of *A3B* is higher than of *A3A*, *A3B* RNA-seq reads misalign to highly homologous exons 3-5 of *A3A* (see **part I**) and incorrectly represent *A3A* expression.

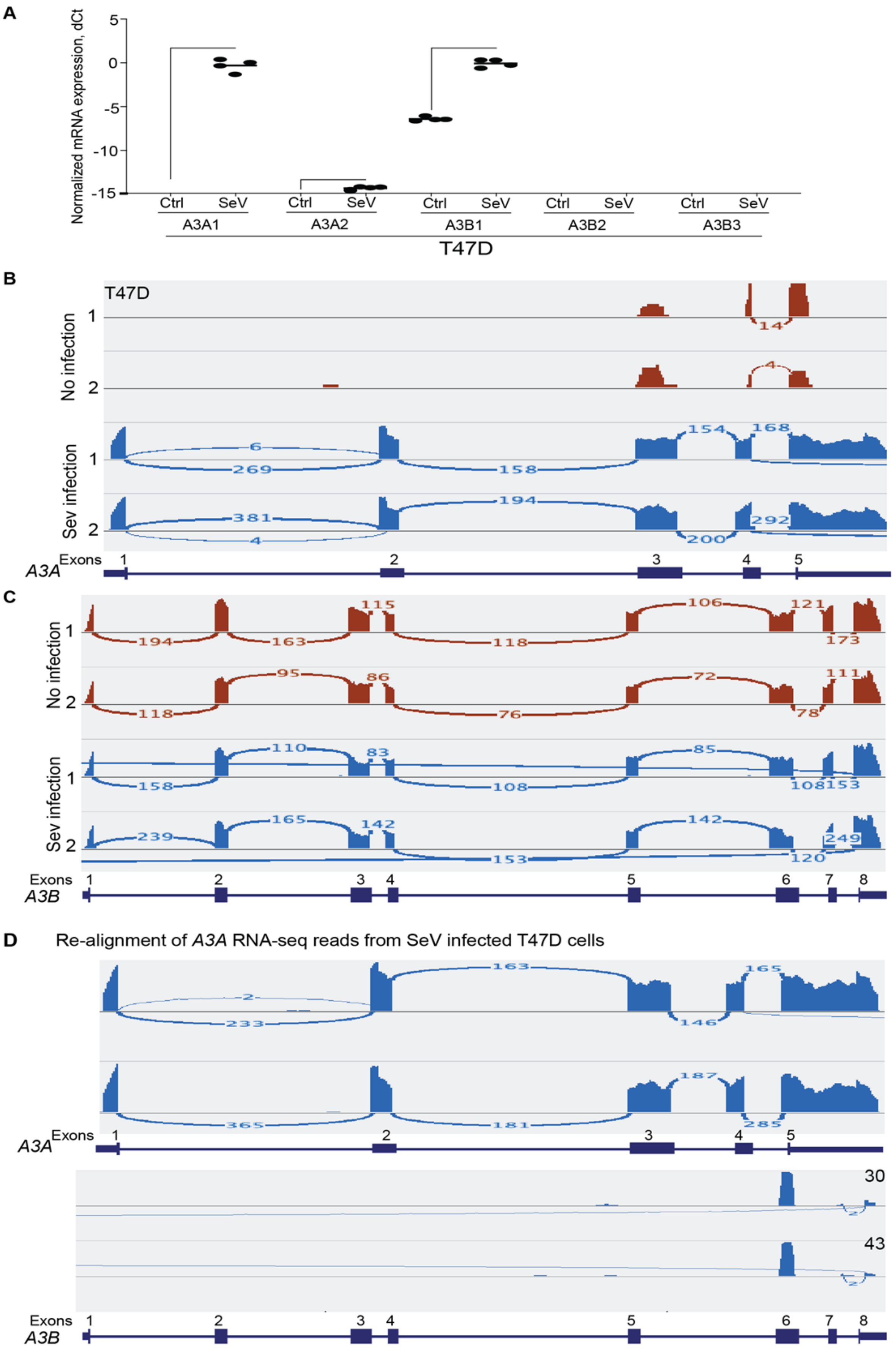

A
Normalized mRNA expression, dCt
5
0
-5
-10
-15
Ctrl
SeV
A3A1
A3A2
A3B1
A3B2
A3B3
T47D
B
T47D
No infection
Sev infection
Exons
A3A
C
A3B
D
Re-alignment of A3A RNA-seq reads from SeV infected T47D cells
30
43

**III). Due to high sequence homology, *A3A* RNA-seq reads misalign to *A3B***

**A**) qRT-PCR analysis of the *A3A* and *A3B* isoforms in a breast cancer cell line T47D in uninfected (control) conditions and in cells infected with Sendai virus (SeV), all in duplicates [7]. After SeV infection, expression of *A3A1* was strongly induced, while *A3B1* was moderately upregulated from baseline levels; **B** and **C**) IGV sashimi plots showing RNA-seq expression profiles of *A3A* and *A3B* in corresponding samples, with *A3A* expression being contaminated by the misaligned *A3B1* reads (see **part II** for details); **D**) Re-alignment of extracted *A3A* reads from SeV-infected T47D cells to the reference genome (hg38) shows misalignment of *A3A1* RNA-seq reads to *A3B* (mostly to exon 6). When expression of *A3A* is higher than of *A3B*, *A3A* RNA-seq reads misalign to highly homologous exon 6 of *A3B* (see **part I**) and incorrectly represent *A3B* expression.

**Note S2. Evaluation of the available antibodies for the detection of A3A and A3B protein isoforms**

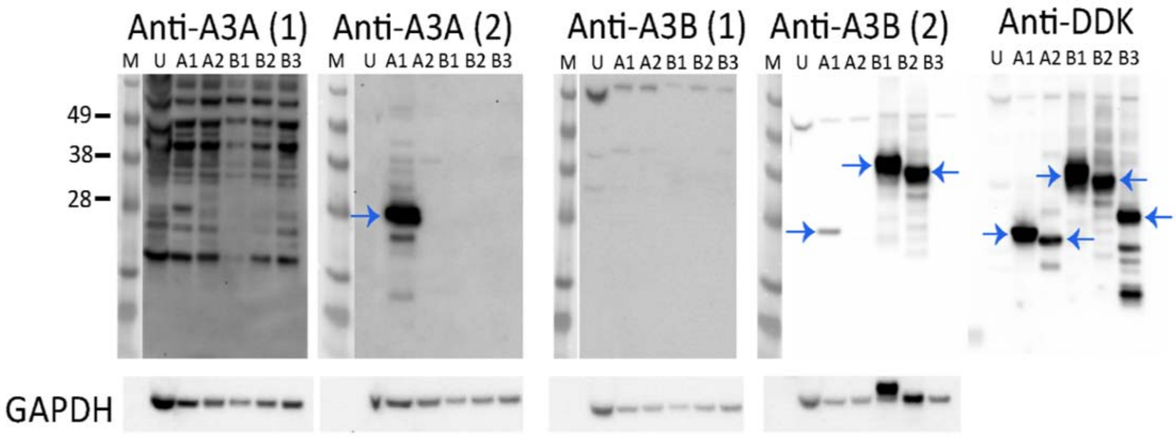


Expression constructs for recombinant Myc-DDK-tagged A3A and A3B protein isoforms (A3A1, A3A2, A3B1, A3B2, and A3B3) were overexpressed in HEK293T cells for 24 hrs. Whole-cell protein lysates of transfected and untransfected (U) cells were resolved on 4-12% SDS PAGE gels. Protein expression was analyzed by Western blots with antibodies described in the table; arrows mark bands considered to be specific.

| Ab | ID | Company | Dilution | Comment |
|---|---|---|---|---|
| A3A (1) | orb19763 | Biorbyt | 1:500 | Not specific |
| A3A (2) | HPA043237 | Millipore Sigma | 1:500 | Detects only A3A1 |
| A3B (1) | ab191695 | Abcam | 1:500 | No detection |
| A3B (2) | ab184990 | Abcam | 1: 500 | Detects A3B1 and A3B2 but not A3B3. Some weak detection of A3A2 as well |
| Anti-DDK | MAB3118 | Millipore Sigma | 1:1000 | Detects all Myc-DDK-tagged proteins |
| Anti-SF3B1 | ab172634 | Abcam | 1:1000 | Detects SF3B1 |
| GAPDH | ab9485 | Abcam | 1:1000 | Loading control |
| mouse-IgGκ BP-HRP | sc-516102 | Santa Cruz Biotechnology | 1:5000 | Secondary ab |
| Anti-goat IgG-HRP | sc-2304 | Santa Cruz Biotechnology | 1:5000 | Secondary ab |
| Anti-rabbit IgG-HRP | 7074 | Cell signaling | 1:5000 | Secondary ab |

GAPDH blot with anti-A3B (2) antibody shows incomplete stripping of the original anti-A3B blot, which was not possible to exclude due to the close size of these bands.

**Note S3. Purification/enrichment of recombinant A3A and A3B protein isoforms**

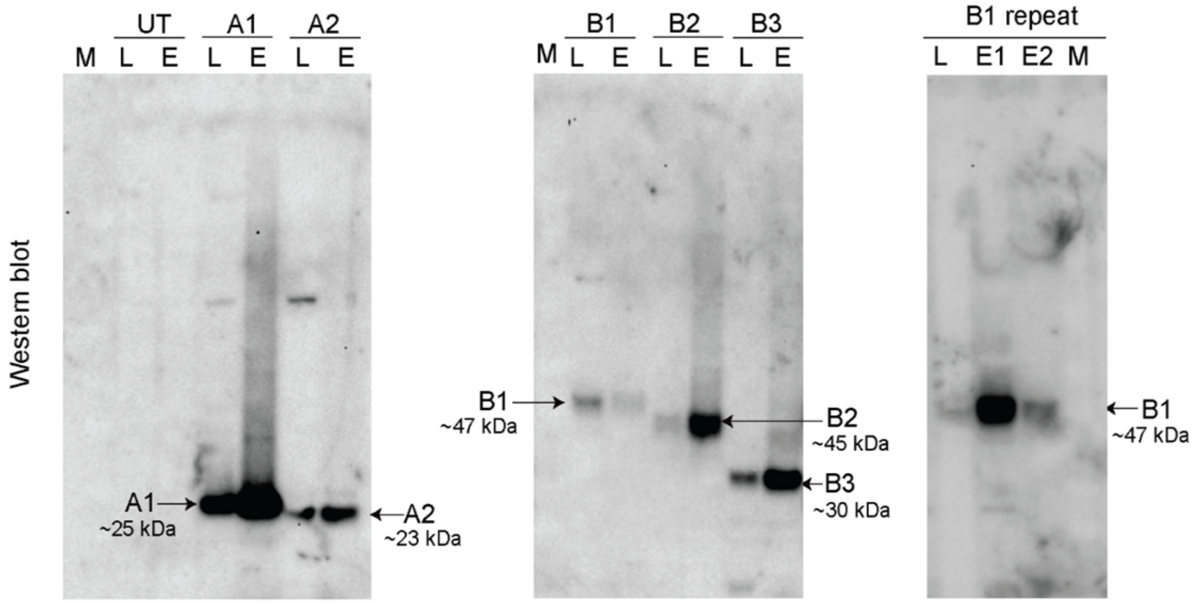


Recombinant Myc-DDK tagged A3A and A3B protein isoforms were purified as described in Materials and Methods. Semi-purified proteins eluted with Myc peptide were resolved on 4-12 % SDS PAGE gels and analyzed by Western blots with an anti-DDK antibody. Each eluate (E) from cells transfected with plasmids for Myc-DDK A3A and A3B isoforms or untransfected (UT) cells was compared with the corresponding cell lysates (L) to determine enrichment of the specific proteins.

#### Note S4. Isoform-specific TaqMan expression assays for *A3B* transcripts

| A. Sequences for primers and probes for custom TaqMan expression assays | | | |
|---|---|---|---|
| **Assay** | **Forward Primer** | **Reverse Primer** | **Probe, MGB (FAM-dye-labeled)** |
| **A3B1** | ACCAGCACATGGGCTTTCTATG | AAAGAAGGAACCAGGTCCAAGAAG | CGAGGCTAAGAATCT |
| **A3B2** | GCTTTCTATGCAACGAGTTGGA | AGGAGAAGCAGGGGCTCC | CAGATCTACAGGGTCACTT |
| **A3B3** | CGCTAAAGGAGATTCTCAGGCTA | CAGGTCCAAGAAGCGCAGC | CTCTGTGGCTTTTACGGC |
| **A3B1&2 for NMD** | TGCTGGGAAAACTTTGTGTACAAT | ATGTGTCTGGATCCATCAGGTATCT | ATTCATGCCTTGGTACAAA |

| **B. Characteristics of custom TaqMan expression assays** | | | | | | | |
|---|---|---|---|---|---|---|---|
| **Assay** | **Plasmid** | **A3A1** | **A3A2** | **A3B1** | **A3B2** | **A3B3** | **Comments** |
| **A3B1** | $R^2$ | 0.70 | 0.97 | **0.99** | 0.94 | 0.70 | Most specific for A3B1 |
| | Ct value | 32.52 | 27.84 | **16.49** | 36.05 | 34.67 | |
| **A3B2** | $R^2$ | 0.79 | 0.77 | 0.86 | **0.99** | 0.99 | Most specific for A3B2 |
| | Ct value | 36.93 | 39.15 | 36.11 | **18.47** | 26.19 | |
| **A3B3** | $R^2$ | 0.98 | 0.52 | 0.98 | 0.36 | **0.99** | Most specific for A3B3 |
| | Ct value | 39.13 | 37.94 | 30.63 | 37.23 | **15.89** | |
| **A3B1&2** | $R^2$ | 0.42 | 0.38 | **0.99** | **0.99** | 0.99 | Most specific for A3B1 and A3B2 |
| | Ct value | 33.34 | 33.40 | **18.19** | **18.73** | 26.47 | |

Expression assays were designed using Primer Express 3.0.1 software (ThermoFisher). The assays were tested using eight 4-fold linear dilutions (from 0.06 to 1000 ng/uL per reaction) of isoform-specific plasmids; $R^2$ coefficients correspond to dilution experiments. Results (Ct values) are shown for 1 ng/uL concentration of all plasmids. Based on analysis with UCSC tools (BLAT and *in silico* PCR), assays are most specific to the designated isoforms. Assays for *A3B1*, *A3B2,* and *A3B3* can also detect other isoforms, but with much lower efficiency.
